# Poly(ADP-ribose) glycohydrolase promotes formation and homology-directed repair of meiotic DNA double-strand breaks independent of its catalytic activity

**DOI:** 10.1101/2020.03.12.988840

**Authors:** Eva Janisiw, Marilina Raices, Fabiola Balmir, Luis Paulin Paz, Antoine Baudrimont, Arndt von Haeseler, Judith L. Yanowitz, Verena Jantsch, Nicola Silva

## Abstract

Poly(ADP-ribosyl)ation is a reversible post-translational modification synthetized by ADP-ribose transferases and removed by poly(ADP-ribose) glycohydrolase (PARG), which plays important roles in DNA damage repair. While well-studied in somatic tissues, much less is known about poly(ADP-ribosyl)ation in the germline, where DNA double-strand breaks are introduced by a regulated program and repaired by crossover recombination to establish a tether between homologous chromosomes. The interaction between the parental chromosomes is facilitated by meiotic specific adaptation of the chromosome axes and cohesins, and reinforced by the synaptonemal complex. Here, we uncover an unexpected role for PARG in promoting the induction of meiotic DNA breaks and their homologous recombination-mediated repair in *Caenorhabditis elegans*. PARG-1/PARG interacts with both axial and central elements of the synaptonemal complex, REC-8/Rec8 and the MRN/X complex. PARG-1 shapes the recombination landscape and reinforces the tightly regulated control of crossover numbers without requiring its catalytic activity. We unravel roles in regulating meiosis, beyond its enzymatic activity in poly(ADP-ribose) catabolism.

## Introduction

Poly(ADP-ribosyl)ation (PARylation) is an essential post-translational modification involved in chromatin dynamics, transcriptional regulation, apoptosis, and DNA repair (Koh et al., 2004; Menissier de Murcia et al., 2003). PARylation is controlled by the opposing activities of PAR polymerases, PARP1 and PARP2 (PARPs), and PAR glycohydrolase (PARG) (O’Sullivan et al., 2019; Slade, 2019). The activities of PARPs are crucial for an efficient DNA damage response, as loss of PARP1 or PARP2 leads to hypersensitivity to genotoxic stress and impaired spermatogenesis, while the combined deficiencies of PARP1 and PARP2 cause embryonic lethality (Dantzer et al., 2006; Menissier de Murcia et al., 2003). Likewise, the PARG knock-out is embryonic lethal and depleted cells become sensitive to ionizing radiation (IR) and show aberrant mitotic progression (Ame et al., 2009; Koh et al., 2004). Since PARP1/2 double mutants or PARG knock-outs are embryonic lethal in mouse and no orthologs are present in yeast, our understanding of the roles of PARylation during germ line development has been limited. *C. elegans parg-1/PARG* null mutants are viable and fertile (Byrne et al., 2016; St-Laurent et al., 2007), allowing us to analyze their function(s) during gametogenesis. It has been previously shown that *parp-1/-2* and *parg-1* mutants display hypersensitivity to IR exposure (Dequen et al., 2005; Gagnon et al., 2002; St-Laurent et al., 2007) however their roles during gametogenesis have remained poorly investigated.

In sexually reproducing species, preservation of ploidy across generations relies on meiosis, a specialized cell division program which promotes the generation of haploid gametes (Zickler and Kleckner, 1999, 2015). The formation of crossovers (CO) is essential for faithful chromosome segregation into the gametes (Cao et al., 1990; Sun et al., 1989). Connected parental homologous chromosomes (also called bivalents) can cytologically be detected in diakinesis nuclei and are thus a readout for the success of the CO establishment. COs arise by the generation and homologous recombination-mediated repair of programmed DNA double-strand breaks (DSB) effectuated by the evolutionarily conserved topoisomerase VI-like protein Spo11 (Keeney et al., 1997). The activity of Spo11 is tightly regulated to ensure the correct timing, placement, and number of DSBs/COs along meiotic chromosome axes. In *C. elegans*, several factors involved in promoting meiotic DSBs have been identified, and those include MRE-11, HIM-5, HIM-17, DSB-1, DSB-2 and XND-1 (Chin and Villeneuve, 2001; Meneely et al., 2012; Reddy and Villeneuve, 2004; Rosu et al., 2013; Stamper et al., 2013; Wagner et al., 2010). Of these, XND-1 and HIM-17 are known to also influence germline chromatin structure (Reddy and Villeneuve, 2004; Wagner et al., 2010). DSB-1 and DSB-2 appear to have roles in maintaining DSB competency throughout early pachytene (Rosu et al., 2013; Stamper et al., 2013). MRE-11 functions both in DSB formation and immediately downstream in end resection (Chin and Villeneuve, 2001); HIM-5 and DSB-2 have also recently been shown to couple DSB formation with HR-mediate repair (Macaisne et al., 2018).

The distribution and number of DSBs and COs also undergoes multiple levels of regulation. In all organisms studied, the number of DSBs exceeds the number of COs, with ratios reaching 10:1 in some cases (Serrentino and Borde, 2012). The excess DSBs use HR-like mechanisms to be repaired with high fidelity, with repair intermediates shunted into non-CO (NCO) outcomes. Importantly, a robust inter-homolog repair bias ensures formation of the obligate CO in the germ cells, which in *C. elegans* occurs even under subthreshold levels of DSBs (Meneely et al., 2012; Rosu et al., 2011; Yokoo et al., 2012). CO interference (Youds et al., 2010; Zickler and Kleckner, 1999) describes the phenomenon whereby CO-committed intermediates influence nearby DSBs to be repaired as NCOs, ensuring that COs are well-spaced across the genome. In *C. elegans*, CO interference is nearly complete, as each chromosome pair receives, in most cases, only one CO (Hillers and Villeneuve, 2003). On the autosomes of the worm, COs occur preferentially on the arms of the chromosomes, away from the gene-rich region in the center of the chromosomes; on the heterochromatic-like X chromosome, there is not gene cluster and COs are more evenly dispersed (Barnes et al., 1995).

While CO interference explains much about CO distribution in most organisms, some COs are known to arise from an interference-independent pathway. The COs generated through interference-dependent (Class I) and interference-independent mechanisms (Class II) have distrinct genetic requirement, driven by MutS-MutL and Mus81 homologs respectively (de los Santos et al., 2003). Genetic evidence suggests that, in *C. elegans*, only Class I COs are present (Kelly et al., 2000; Zalevsky et al., 1999). Nevertheless mutants displaying interference-insensitive COs have been reported (Tsai et al., 2008; Youds et al., 2010), however, these are still dependent on the canonical MSH-5/COSA-1-mediated CO pathway and they can be detected by genetic measurements of recombination (Yokoo et al., 2012).

CO-repair takes place in the context of the synaptonemal complex (SC), a tripartite proteinaceous structure composed of axial and central elements, arranged as a protein-zipper between each pair of homologs. The SC maintains homolog associations and facilitates inter-homolog exchange of DNA during repair (Colaiacovo et al., 2003). Cross-talk between the SC and COs is essential for modulating recombination. Incomplete synapsis dramatically weakens CO interference and additional COs *per* chromosome can be observed (Libuda et al., 2013; Rosu et al., 2011). Conversely, reduced, but not absent, recombination levels causes premature desynapsis of the chromosome pairs that fail to establish a CO (Machovina et al., 2016; Pattabiraman et al., 2017; Wagner et al., 2010).

Chromosome axis components, which in *C. elegans* include the HORMA-domain proteins HTP-3, HTP-1/2, and HIM-3 (Goodyer et al., 2008; Martinez-Perez and Villeneuve, 2005; Zetka et al., 1999), influence both the abundance of DSBs and the regulation of their repair.

In this study, we show an unexpected involvement of PARG-1 in promoting DSB and CO formation. We show that PARG-1 functions independently of the known DSB initiation factors and that it cooperates with HIM-5 to regulate global crossover numbers. PARG-1 is detected throughout the germ line and undergoes a progressive recruitment along synapsed chromosomes, culminating in the retraction to the short arm of the bivalent and enrichment at the putative CO sites. In the absence of PARG-1 function, we observe an accumulation of PAR on the meiotic chromosomes, which is suppressed by abrogation of PARP-2 function.

We report the association of PARG-1 with numerous key proteins composing the meiosis-specific structure of the SC both by cytological and biochemical analysis. Surprisingly, we found that PARG-1 loading, rather than its catalytic activity, is essential to exert its function during meiosis. Our data strongly suggest that PARG has scaffolding properties which are important for the fine-tuning of meiotic recombination events.

## Results

### PARG-1 is the main poly(ADP-ribose) glycohydrolase in the *C. elegans* germ line

The *C. elegans* genome encodes two orthologs of mammalian PARG, PARG-1 and PARG-2 (Bae et al., 2019; Byrne et al., 2016; St-Laurent et al., 2007). Both mutants are hypersensitive to IR exposure and more recently it was shown that *parg-2* is involved in the regulation of HR-dependent repair of ectopic DSBs by influencing the extent of resection upon IR (Bae et al., 2019). To explore possible functional links or redundancies between *parg-1* and *parg-2*, we used CRISPR to engineer *parg-2* null mutations in both the wild type (WT) and *parg-1(gk120)* deletion mutant backgrounds (Fig.1A). In contrast to mammalian PARG, *C. elegans parg-1* and *parg-2* are largely dispensable for viability (Fig. 1B). However, abrogation of *parg-1*, but not *parg-2* function, led to increased levels of embryonic lethality and segregation of males (which arise from X chromosome nondisjunction (Hodgkin et al., 1979). Screening of *parg-1 parg-2* double mutants did not reveal synthetic phenotypes but recapitulated the *parg-1* phenotypic features, indicating that *parg-2* does not exert prominent roles in an otherwise wild type background and cannot compensate the lack of *parg-1* function (fig. 1B).

**Figure 1.**
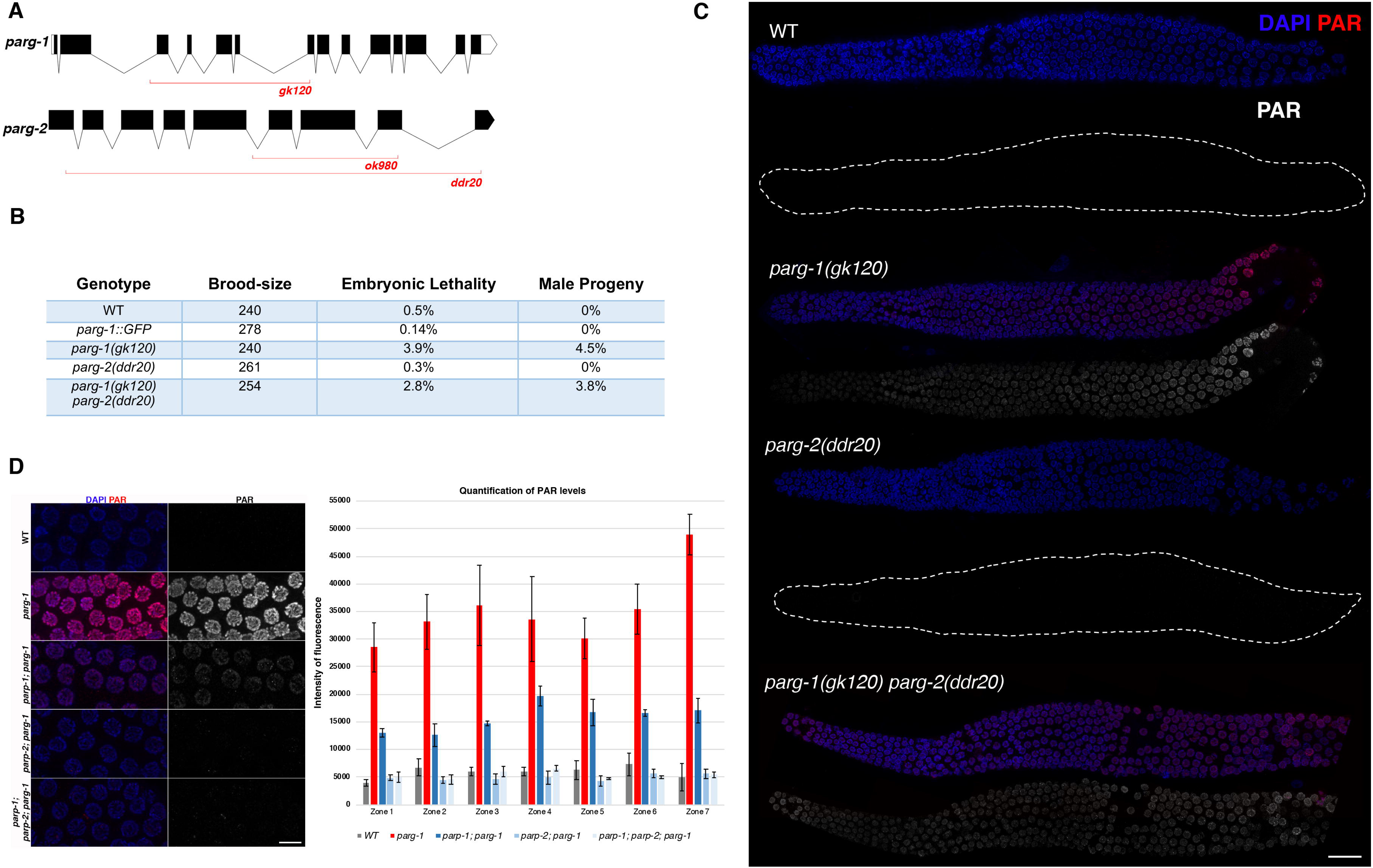
PARG-1 is the main ADP-ribose glycohydrolase in *C. elegans.* **(A)** Schematic representation of *parg-1* and *parg-2* genetic loci. *parg-1* is predicted to encode numerous isoforms and for simplicity only the transcript encoding isoform A is shown. Red lines delineate the position of deletions in the indicated mutant alleles. **(B)** Screening of brood-size, embryonic lethality and segregation of males in indicated genetic backgrounds. Number of embryos scored: WT (1196), *parg-1::GFP (*1389), *parg-1(gk120)* (1200), *parg-2(ddr20)* (1304) and *parg-1(gk120) parg-2(ddr20)* (1271). **(C)** Representative images of whole-mount gonads from indicated genotypes showing detection of PAR by immunofluorescence. Scale bar 20 μm. **(D)** Left: representative pictures of late pachytene nuclei from indicated genotypes showing *parp-1-* and *parp-2*-dependent accumulation of PAR in *parg-1* mutants. Gonads were divided into 7 zones, encompassing the region from the distal tip cell to diplotene. Scale bar 5 μm. Right: quantification of PAR detected by immunofluorescence. Chart reports average of fluorescence intensity from at least two gonads/genotype. Error bars indicate standard deviation.

To confirm a role of PARG-1 and PARG-2 in PAR catabolism, we investigated PAR accumulation in the mutant animals. Because PAR undergoes a rapid turnover, PAR cannot be detected in wild-type germ lines (Fig. 1C). By contrast, we detected PAR at all stages of meiotic prophase I in *parg-1* mutants. Since PAR accumulation was neither seen in *parg-2* mutants nor further enhanced in *parg-1 parg-2* (Fig.1C), we infer that PARG-1 is the major PAR glyohydrolase in the worm germ line.

Removal of the PAR polymerases *parp-1/-2*, suppressed accumulation of PAR in *parg-1* mutant germ cells (Fig. 1D). Interestingly, we found that while abrogation of *parp-1* function reduced detectable levels to roughly 30%, lack of *parp-2* alone was sufficient to bring PAR staining to background levels. Since both *parp-1(ddr31)* and *parp-2(ok344)* mutant alleles are null, this data suggests that PARP-2 is mainly responsible for the synthesis of PAR during *C. elegans* meiotic prophase I.

Since PAR accumulates at sites of DNA damage in somatic cells (Kaufmann et al., 2017; Mortusewicz et al., 2011), we asked whether the accumulation of PAR in meiotic prophase nuclei was dependent on the formation of meiotic DSBs. Surprisingly, we found that in the gonads of *parg-1 spo-11* double mutants, in which no programmed DSBs are made, PAR still localized within prophase I nuclei (Fig. S1), indicating that PAR synthesis occurs independently of physiological DNA damage during gametogenesis.

### PARG-1 forms protein complexes with SC components and its localization requires chromosome axes

To detect PARG-1, we raised a *C. elegans*-specific anti-PARG-1 monoclonal antibody which we used in western blot analysis (Fig. 2A). This antibody confirmed that *parg-1(gk120)* is a null allele. We find expression of PARG-1 in all subcellular compartments in wild-type animals (Fig. 2A), as similarly observed in mammalian mitotic cells (Meyer-Ficca et al., 2004; Ohashi et al., 2003; Winstall et al., 1999). Since localization of PARG is not known in meiocytes, we employed CRISPR to tag the 3’ end of the endogenous *parg-1* locus with a GFP-tag. We assessed the functionality of the fusion protein by monitoring PAR accumulation in the gonad, embryonic lethality, and male progeny, none of which showed any differences compared to WT, indicating that PARG-1::GFP is catalytically active and fully functional (Fig. 1B-C and S1). Moreover, western blot analysis employing either anti-PARG-1 or anti-GFP antibodies on fractionated extracts from *parg-1::GFP* worms revealed identical expression as seen with untagged PARG-1 (Fig. 2B), further confirming that the GFP-tag did not affect PARG-1 stability or expression.

**Figure 2.**
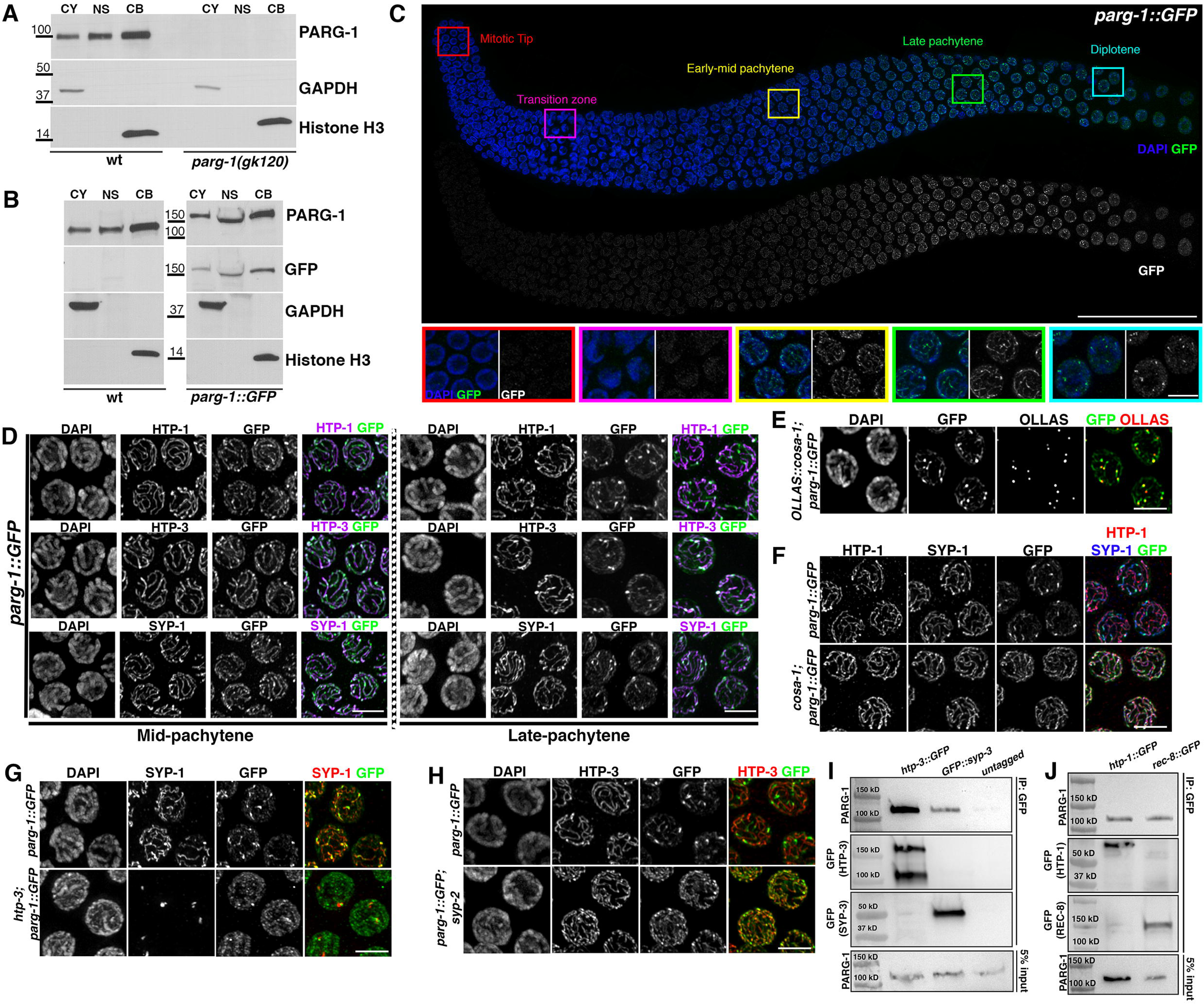
PARG-1 interacts with SC components and requires chromosome axes for proper localization. **(A)** Western blot analysis of fractionated extracts detects PARG-1 in all subcellular compartments with enrichment in the chromatin-bound fraction. CY= cytosol, NS= nuclear soluble, CB= chromatin bound. GAPDH was used as a loading control of the cytosolic fraction and Histone H3 for the chromatin bound fraction. **(B)** Western blot analysis of fractionated extracts showing similar expression of GFP-tagged and untagged PARG-1. **(C)** Top: PARG-1::GFP localization in a wild-type gonad. Scale bar 30 μm. Bottom: enlarged insets showing dynamic localization of PARG-1::GFP in different stages of meiotic prophase I. Scale bar 5 μm. **(D)** Mid-and late-pachytene nuclei of *parg-1::GFP* co-stained for lateral (HTP-1 and −3) and central component (SYP-1) of the SC. Scale bar 5 μm. **(E)** Late pachytene nuclei showing overlapping localization of PARG-1::GFP with OLLAS::COSA-1. Scale bar 5 μm. **(F)** Late pachytene nuclei stained for HTP-1, SYP-1 and GFP showing localization of PARG-1 along chromosomes. In *cosa-1* mutants redistribution to the short arm of the bivalent is absent. Scale bar 5 μm. **(G)** Impaired axes formation in *htp-3* mutants prevents PARG-1::GFP localization. Scale bar 5 μm. **(H)** PARG-1::GFP associates with HTP-3 in late-pachytene nuclei in absence of synapsis. Scale bar 5 μm. **(I)** Western blot analysis of endogenous PARG-1 on GFP pull downs performed in *htp-3::GFP* and *GFP::syp-3* strains. Wild-type worms were used as the untagged negative control. **(J)** Western blot analysis of endogenous PARG-1 on GFP pull downs performed in *htp-1::GFP* and *rec-8:: GFP* strains.

Immunofluorescence analyses showed that PARG-1::GFP is first detected in pre-meiotic and leptotene/zygotene nuclei and then became progressively enriched along chromosomes throughout pachytene (Fig. 2C). Co-staining with axial proteins HTP-1/HTP-3 and the central SC component SYP-1 (Goodyer et al., 2008; MacQueen et al., 2002; Martinez-Perez and Villeneuve, 2005) revealed recruitment of PARG-1::GFP onto synapsed chromosomes and its retraction in late pachytene cells to the short arm of the bivalent (Fig. 2D), a chromosomal subdomain formed in response to CO formation. The short arm of the bivalent harbors the chiasma and the central elements of the SC (de Carvalho et al., 2008; Martinez-Perez et al., 2008). Overlapping localization of PARG-1::GFP with both the CO promoting factor COSA-1 and SYP-1 (Fig. 2E) confirmed recruitment of PARG-1 to this chromosomal subdomain, similar to other SC central elements components (Bhalla et al., 2008; Janisiw et al., 2018; Jantsch et al., 2004; Li et al., 2018; MacQueen et al., 2002). In CO-defective *cosa-1* mutant animals, we observed that the initial loading of PARG-1::GFP to the SC was unaffected, but no retraction was observed, confirming that the relocalization of PARG-1 is dependent on bivalent formation (Fig. 2F).

Based on its localization to the SC, we tested whether PARG-1::GFP loading was dependent on chromosome axis or synapsis establishment. Loss of *htp-3*, encoding a HORMA domain-containing protein essential for axis morphogenesis (Goodyer et al., 2008), disrupted PARG-1::GFP localization, resulting in nucleoplasmic accumulation and occasional association with SYP-1-containing polycomplexes (Fig. 2G). By contrast, PARG-1::GFP exhibited linear staining along the chromosome axes in synapsis-deficient *syp-2* mutants (Fig. 2H), where only axial elements are loaded onto the chromosomes (Colaiacovo et al., 2003; Goodyer et al., 2008; Martinez-Perez and Villeneuve, 2005). Thus, we conclude that PARG-1 is recruited to the SC in an HTP-3-dependent manner and its localization changes in response to CO-mediated chromosome remodeling.

Since PARG-1 localizes to chromosome axes and requires HTP-3 for loading, we wondered whether these factors formed protein complexes *in vivo*. To test for their possible association, we performed immunoprecipitation assays by pulling down HTP-3::GFP (Paix et al., 2015) and proceeded with western blot analysis to detect PARG-1. Robust interaction between HTP-3::GFP and PARG-1 was observed (Fig. 2I). Further, to assess whether PARG-1 establishes physical interactions with additional chromosome axis components as well, we also performed co-immunoprecipitation experiments pulling down HTP-1::GFP and REC-8::GFP, (Crawley et al., 2016; Silva et al., 2014). Western blot showed that PARG-1 co-immunoprecipitated with both HTP-1 and REC-8 (Fig. 2J). Extending these analysis to the central elements of the SC component, we found that PARG-1 could be pulled down with GFP::SYP-3 (Rog and Dernburg, 2015) (Fig. 2I). Together with our localization studies, these physical interactions indicate that PARG-1 is an intrinsic component of the SC.

### Loss of *parg-1* suppresses chromosome abnormalities arising from impaired DSB resection

Given PARG-1 recruitment along the SC and enrichment at the presumptive CO sites, we sought to investigate whether synapsis and CO formation are impaired in *parg-1* mutants. Using antibodies directed against HTP-3 and SYP-1 to monitor the establishment of the SC, we observed no difference between the wild type and *parg-1* mutants (Fig. S2A). DAPI-staining of diakinesis nuclei revealed the correct complement of six bodies as in wild-type worms (Fig. S2B). Thus, we infer that *parg-1* is dispensable for synapsis and CO formation.

We next addressed whether loss of *parg-1* would impact the formation and processing of recombination intermediates by analyzing the dynamic behaviour of the recombinase RAD-51, which forms discrete chromatin-associated foci with a distinct kinetics of appearance and disappearance (Alpi et al., 2003; Colaiacovo et al., 2003), (Fig. 3A-B). While in wild-type worms we see a progressive increase of RAD-51, peaking in early-mid pachytene (zone 3) and disappearing by late pachytene (zone 6), in *parg-1* mutants, we observed the delayed formation of RAD-51 foci with progressive accumulation at the pachytene stage. RAD-51 foci formation was entirely suppressed by SPO-11 removal, suggesting specific abnormalities in the induction and/or processing of meiotic DSBs rather than spontaneous or unscheduled damage arising during mitotic replication (Fig. 3A-B).

**Figure 3.**
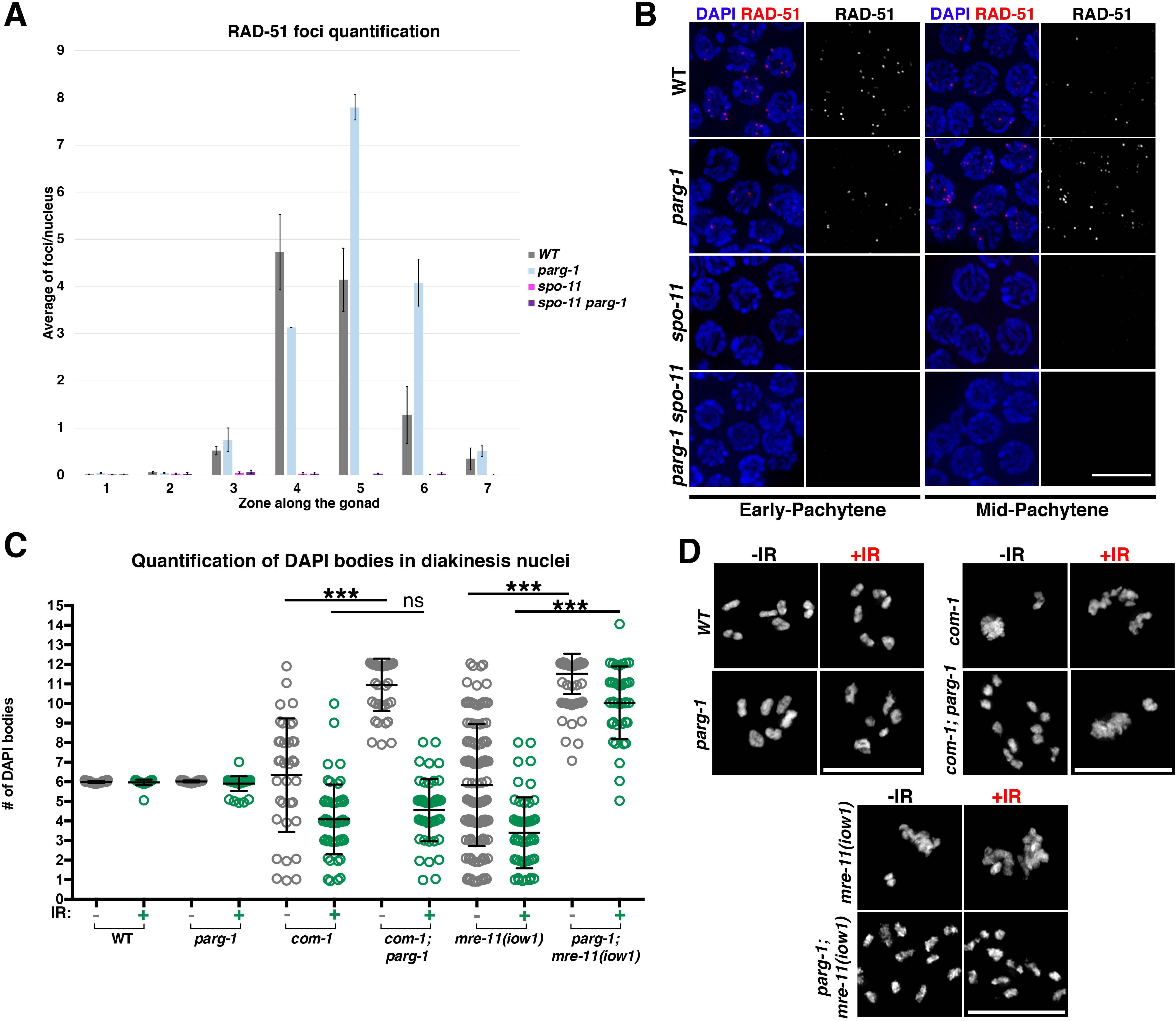
Elimination of *parg-1* function suppresses chromosome abnormalities in resection-defective mutants. **(A)** *parg-1* mutants display SPO-11-dependent accumulation of RAD-51 foci. Error bars represent S.E.M. **(B)** Representative examples of cells at different stages of the same genotypes analyzed in A. Scale bar 5 μm. **(C)** Analysis of diakinesis nuclei in different genotypes before and 27h after exposure to IR. Error bars represent standard deviation. Statistical analysis was performed with non-parametric Mann-Whitney test. *** indicates *p<0.0001* and *ns* indicates statistically non-significant differences. **(D)** Representative images of diakinesis nuclei of the same genotypes scored in C. Scale bar 5 μm.

Since RAD-51 foci appeared with delayed kinetics, we wanted to investigate whether PARG-1 might have a role in the regulation of DSB formation. Since tools to directly quantify meiotic DSBs are presently not available in *C. elegans*, we took advantage of a genetic epistasis analysis to determine if *parg-1* has a role in DSB formation. In diakinesis nuclei, DSB resection-defective mutants, such as *com-1/CtIP/Sae2* and *mre-11(iow1)/Mre11*, display massive chromatin clumps and occasional chromosome fragments that arise from aberrant repair of meiotic DSBs. Accordingly, these clumps and fragments are fully suppressed in the DSB-devoid *spo-11* mutants (Chin and Villeneuve, 2001; Penkner et al., 2007; Yin and Smolikove, 2013). Similarly, in the *com-1; parg-1* and *parg-1; mre-11(iow1)* double mutants, we found that the vast majority of diakinesis nuclei contained twelve intact univalents (Fig. 3C-D). These results are consistent with a role for PARG-1 in DSB induction, but could also reflect a function for *parg-1* in targeting breaks to alternative repair pathways.

To distinguish between these possibilities, we exposed the aforementioned double mutants to gamma irradiation (IR) to ectopically induce DSBs. We reasoned that if *parg-1* mutants were defective solely in DSB induction, the breaks induced by IR should restore the aberrant chromosome morphology typical of *com-1* and *mre-11*. By contrast, if *parg-1* has a role in repair pathway utilization, the IR-induced breaks would still be shunted into an alternative pathway and the appearance of DAPI bodies would remain unchanged after IR exposure. Diakinesis nuclei of irradiated *com-1; parg-1* reverted to the *com-1*-like (chromosome clumping-fusion) phenotype, supporting a putative role for PARG-1 in DSB induction. By contrast, *parg-1; mre-11(iow1)* were indistinguishable from non-irradiated controls, indicating that PARG-1 may also influence DNA repair pathway choice when *mre-11*, but not *com-1*, function is compromised (Fig. 3C-D). Together, these results suggest an involvement of PARG-1 in promoting both the formation and repair of meiotic DSBs.

### PARG-1 augments the formation of meiotic DSBs and interacts with MRE-11

To further explore PARG-1’s putative involvement in promoting DSBs, we tested its ability to genetically interact with mutations that are impaired in DSB induction. We combined the *parg-1(gk120)* deletion with two hypomorphic *him-17* alleles and with a *him-5* null mutation that reduce, but do not completely eliminate, SPO-11-dependent DNA breaks (Meneely et al., 2012; Reddy and Villeneuve, 2004). Consistent with published results, we observed that these single mutants displayed reduced numbers and delayed formation of RAD-51 foci (Meneely et al., 2012; Reddy and Villeneuve, 2004). The number of RAD-51 foci was further diminished in both *parg-1; him-5* and *parg-1; him-17* double mutants (Fig. S3A). The defects in RAD-51 filament formation in *parg-1; him-17* and *parg-1; him-5* double mutants were correlated with defective loading of pro-CO factors such as HA::RMH-1, GFP::MSH-5 and OLLAS::COSA-1 (Fig. 4B-C and S4) as we would expect for mutations that impair DSB formation. Analysis of diakinesis nuclei revealed an extensive lack of chiasmata (Fig 4D) and enhancing embryonic lethality (Fig S3B) in the double mutants as expected from the defects in CO repair. Loading of RMH-1, MSH-5 and COSA-1, as well as bivalent formation, were largely, although not completely, rescued by IR exposure (Fig. 4A-D and S4), further corroborating that the lack of COs was due to impaired DSB formation. Abrogation of PARG-1 function also exacerbated the CO defect observed in both young (day #1) and old (day #2) *dsb-2* mutants (Fig. S3C), which display an age-dependent loss in the proficiency to induce DSBs (Rosu et al., 2013). These results indicate that loss of *parg-1* function impairs a parallel, *him-17-, him-5,* and *dsb-2*-independent pathway for DSB induction.

**Figure 4.**
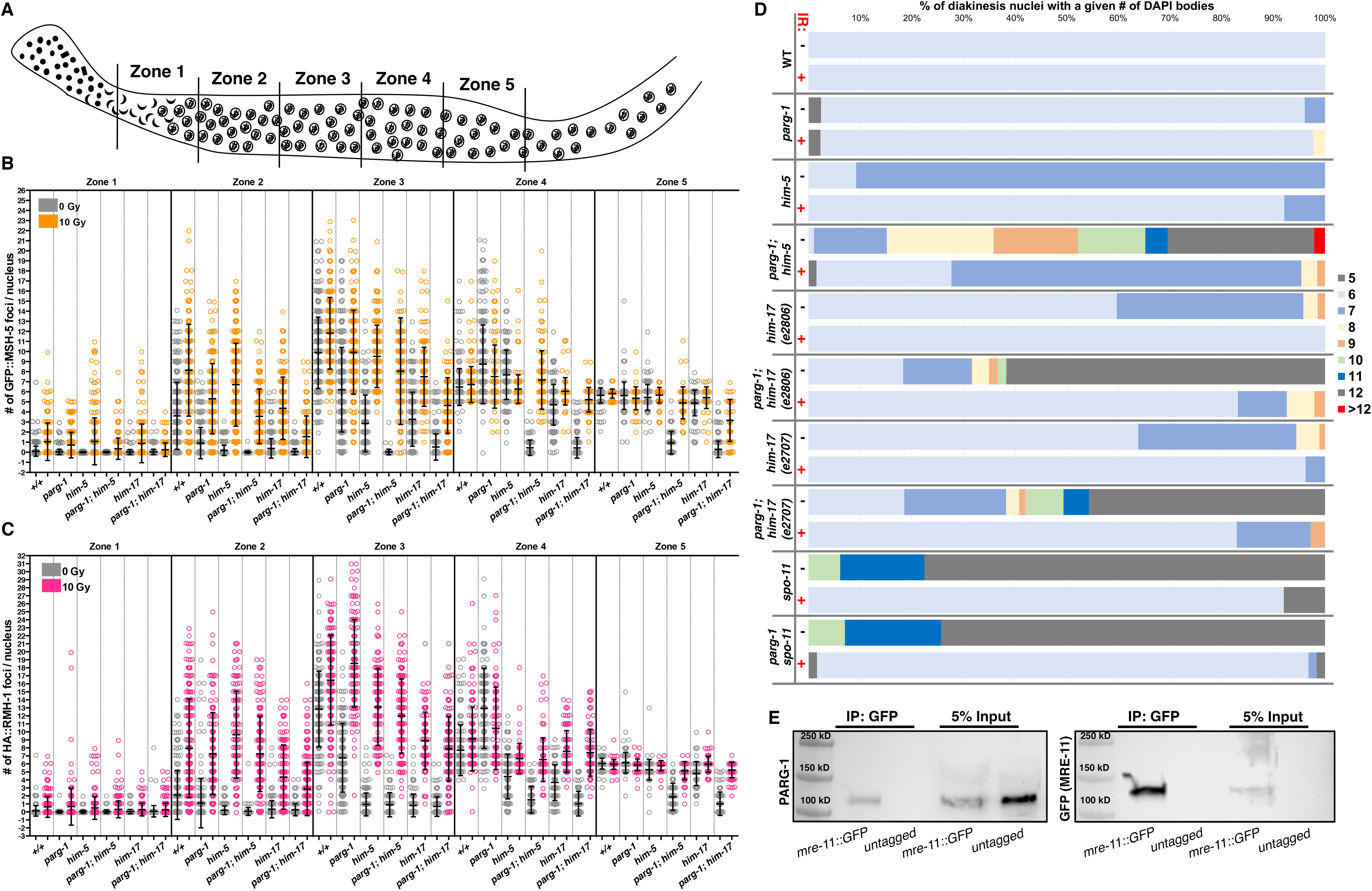
PARG-1 promotes formation of meiotic DSBs and interacts with MRE-11. **(A)** Schematic representation of the gonad divided into five equal regions spanning transition zone throughout late pachytene, employed for MSH-5 and RMH-1 foci quantification. **(B)** Quantification of GFP::MSH-5 foci in the indicated genotypes before and after IR. Error bars indicate standard deviation. **(C)** Quantification of HA::RMH-1 foci in the indicated genotypes before and after IR. **(D)** Quantification of DAPI-bodies of different genotypes before and after IR exposure. **(E)** Western blot analysis of endogenous PARG-1 on GFP pull downs performed in *mre-11::GFP* and untagged wildtype strains (negative control).

To further interrogate PARG-1 function in DSB formation, we next sought to investigate the interplay between PARG-1 and DSB-promoting factors. To this end, we assessed the localization of the pro-DSB factors HIM-5::3xHA, HIM-17::3xHA, DSB-2 and XND-1 in *parg-1* mutants. We observed no gross defects in localization compared to the controls (Fig. S5A-D), which suggests that PARG-1 is not required for the loading of these pro-DSB factors. Conversely, PARG-1::GFP loading appeared normal in *him-5*, *dsb-2*, and *him-17* (null and hypomorph alleles) mutant backgrounds. The only difference compared to wild type is the lack of retraction of PARG-1::GFP to the short arm of the bivalent, which is a consequence of the lack of COs caused by these mutations (Fig. S5E) similar to *cosa-1* mutations (described above). Given the synergistic phenotypes observed in the double mutants and the lack of defects in the loading/expression of DSB-promoting proteins, we conclude that PARG-1 supports formation of DSB *via* alternative pathway(s) to the known pro-DSB factors in *C. elegans*.

It has been previously shown that the axial component HTP-3 promotes the formation of meiotic DSBs in worms possibly through its interaction with the MRN/X complex factor MRE-11, known to be involved in DSB formation (Goodyer et al., 2008; Hayashi et al., 2007; Yin and Smolikove, 2013). Since we already showed an interaction between HTP-3::GFP and PARG-1 (Fig. 2I), we now wanted to address if this extended to an association with MRE-11. Western blot analysis for PARG-1 on GFP pull downs performed with the *mre-11::GFP* transgene (Reichman et al., 2018) also showed coimmunoprecipitation (Fig. 4E). This suggests that the PARG-1-mediated activity in promoting meiotic DSBs may intersect the HTP-3-MRE-11 axis.

### PARG-1 and HIM-5 modulate crossover numbers

While the loading of pro-CO factors was largely rescued in the irradiated *parg-1; him-5* double mutants, over half of the diakinesis nuclei still displayed univalents (Fig. 4D), indicating substantial, yet incomplete, restoration of chiasmata. The dose employed in our irradiation experiments (10 Gy) sufficed to fully elicit bivalent formation in *him-5*, *spo-11*, *parg-1 spo-11* and *spo-11; him-5* (Mateo et al., 2016), (Fig. 4D). Therefore, we conclude that additional CO execution steps are defective in *parg-1; him-5*. Importantly, this phenotype was not observed in *parg-2; him-5* double mutants, in which the number and structure of DAPI bodies resembled *him-5* single mutants both before and after exposure to IR (Fig. S6). We thus confirmed that the recombination defects observed in *parg-1; him-5* are specific to impaired *parg-1* function.

The SC is a dynamic structure that responds to the presence or absence of (as yet unidentified) CO intermediates in the nucleus. When COs are made, they stabilize the SC in *cis*. In genetic backgrounds with reduced DSB induction, such as those described above, the chromosome pairs lacking a CO undergo desynapsis at a late pachytene stage, whereas in mutants that completely lack COs, homologs remain fully synapsed but the SC subunits are more labile (Machovina et al., 2016; Pattabiraman et al., 2017). Given the partial rescue of chiasmata formation in *parg-1; him-5* double mutants after IR and the localization of PARG-1 to the SC, we sought to determine if CO designation and SC dynamics are decoupled by simultaneous loss of both HIM-5 and PARG-1 functions.

In unirradiated *him-5* mutant worms, the sole absence of a CO on chromosome X caused its extensive desynapsis in late pachytene nuclei (Fig. 5A, C), recapitulating previous observations (Machovina et al., 2016). By contrast, in *parg-1; him-5* double mutants, the majority of nuclei showed full synapsis (Fig. 5A-C), in agreement with the fact that de-synapsis is not triggered when CO establishment is fully abrogated (Machovina et al., 2016; Pattabiraman et al., 2017). In support of this interpretation, we show that nuclei containing fully synapsed chromosomes displayed no COSA-1 loading in unirradiated *parg-1; him-5* double mutants (Fig. 5D). Immunostaining for H3K4me2, a histone modification that shows specific enrichment on the autosomes, but not on the X chromosome (Reuben and Lin, 2002), further revealed that the X chromosome was fully synapsed in *parg-1; him-5* doubles, consistent with the lack of a CO and in stark contrast to *him-5* mutants (Fig. 5C).

**Figure 5.**
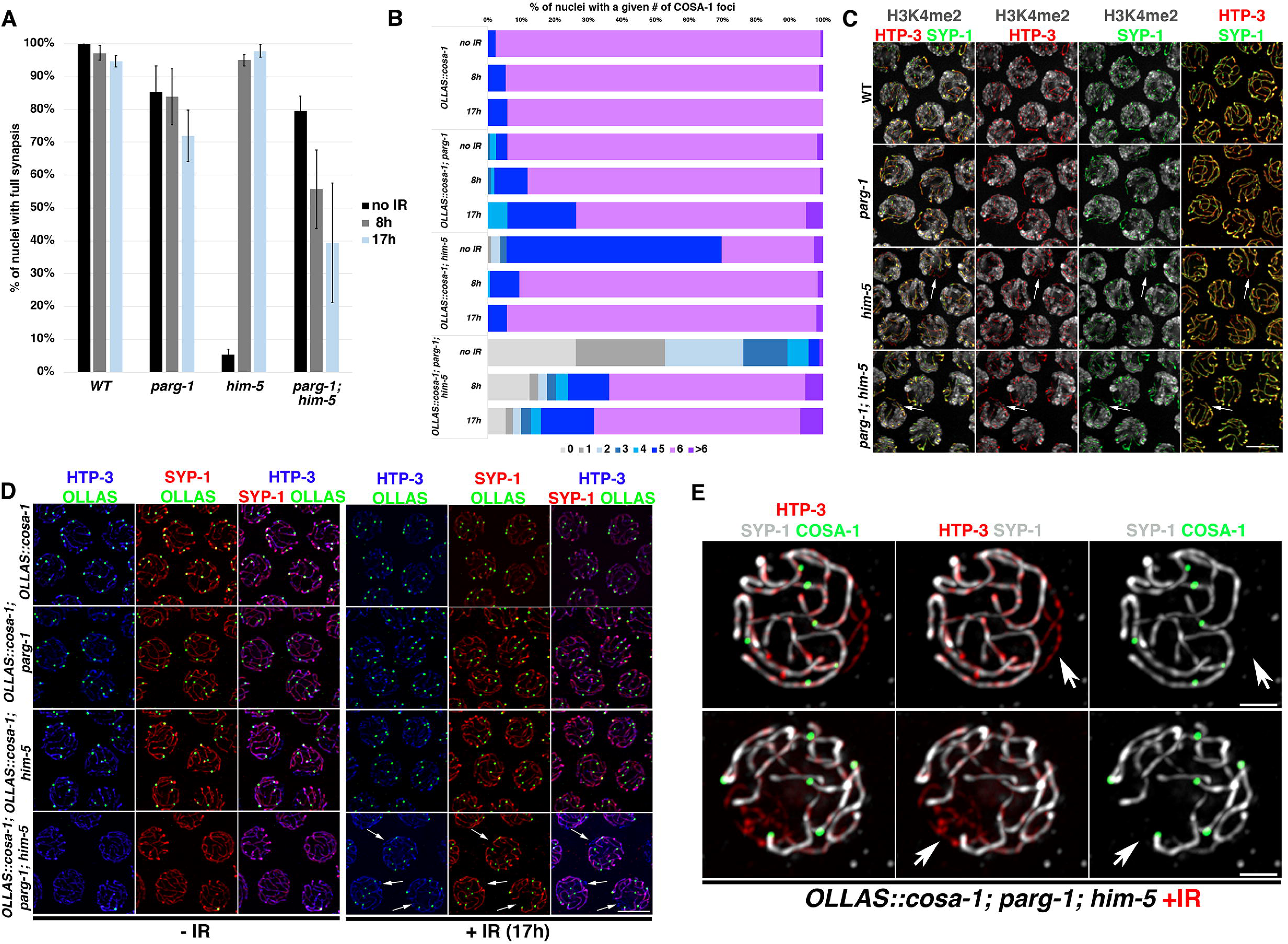
PARG-1 and HIM-5 regulate CO numbers. **(A)** Quantification of synapsis in late pachytene nuclei without IR and at different times after IR exposure, by SYP-1 and HTP-3 co-staining. Only nuclei showing complete colocalization of HTP-3 and SYP-1 were considered fully synapsed. Quantification was performed in the last seven rows of nuclei before entering the diplotene stage. Error bars indicate S.E.M. **(B)** Quantification of OLLAS::COSA-1 foci formation in late pachytene nuclei in the same genotypes and at the same time points as in A. **(C)** Immunostaining of H3K4me2, HTP-3 and SYP-1 to assess chromosome X synapsis in different genotypes. Scale bar 5 μm. **(D)** Co-staining of OLLAS::COSA-1 with SYP-1 and HTP-3 shows de-synapsis associated with lack of CO but normal numbers of COSA-1 foci on the remaining chromatin in *parg-1; him-5* double mutants. Scale bar 5 μm. Arrows indicate examples of unsynapsed regions in nuclei containing six COSA-1 foci. **(E)** High magnification of late pachytene *parg-1; him-5* nuclei after 8h (top) and 17h (bottom) post irradiation of showing normal numbers of COSA-1 foci despite de-synapsis. Arrows indicate desynapsed chromosome regions (presence of HTP-3, absence of SYP-1). Scale bar 1 μm.

We next wanted to address whether SC stabilization and CO formation are coordinated in the *parg-1; him-5* double mutants after irradiation, where six COSA-1 foci were observed (Fig. 5B) but univalents resulted (Fig. 4D). For this analysis, we undertook a time course analysis. In the *him-5* single mutant, 10 Gy of IR is sufficient to both rescue COSA-1 loading and to suppress X chromosome desynapsis as observed both 8 hr and 17 hr post-IR, as shown previously (Machovina et al., 2016). No defects were observed in the *parg-1* single mutants. In *parg-1; him-5*, COSA-1 foci numbers were also largely rescued at 8 hr post-IR and remained steady at 17-hours post-IR (Fig 5B). However, synapsis levels started to decline at 8 hr post-IR and were further reduced at 17-hours after irradiation (Fig. 5A). Strikingly, a substantial number of nuclei exhibited desynapsis, yet they showed the full complement of six COSA-1 foci (8 hr= 52% and 17 hr= 74.3%) (Fig. 5D-E), a situation never described in other meiotic mutants. COSA-1 foci were never associated with unsynapsed chromosome(s) at the observed time points. The fact that these nuclei contain six COSA-1 foci, as in wild-type animals, suggests that some chromosomes might bear additional COSA-1 marked CO events. These results revealed that the global regulation of CO-mediated DNA repair is profoundly perturbed in the absence of PARG-1 and HIM-5 functions.

To further characterize the defects in *parg-1; him-5* mutants, we examined the meiotic progression marker phospho-SUN-1^S8^ (Penkner et al., 2009). In wild-type animals, SUN-1^S8^ is phosphorylated in leptotene/ zygotene and dissipates at mid-pachytene (Woglar et al., 2013). The lack of DSBs or impaired homologous recombination-mediated repair trigger retention of phospho-SUN-1^S8^ at the nuclear envelope until the late pachytene stage (Woglar et al., 2013). In DSB-defective mutants, but not in mutants with impaired recombination (such as *cosa-1*), delayed removal of phospho-SUN-1^S8^ is rescued by exogenous DSB induction (Machovina et al., 2016; Rosu et al., 2013; Stamper et al., 2013; Woglar et al., 2013). Since *parg-1; him-5* double mutants appear to carry defects in both DSB induction and repair, we analyzed phospho-SUN-1^S8^ localization before and after IR exposure to assess whether these phenotypes could be uncoupled by phospho-SUN-1^S8^ dynamics. *parg-1* mutants displayed mild prolongation of phospho-SUN-1^S8^ staining (Fig. S6), consistent with the delayed accumulation of RAD-51 foci (Fig. 3A and S3). *him-5* and *parg-1; him-5* mutants showed comparable, prolonged phospho-SUN-1^S8^ staining under unchallenged growth conditions, consistent with defective DSB induction and recombination. While IR exposure fully suppressed the persistence of phospho-SUN-1^S8^ in the *him-5* as expected, it only mildly suppressed it in *parg-1; him-5* (Fig. S7). The inability of IR to suppress phospho-SUN-1^S8^ accumulation further reinforces the conclusion that lack of both PARG-1 and HIM-5 impairs both meiotic DSB formation and repair.

### PARG-1 shapes the recombination landscape and reinforces CO interference

Given the involvement of *parg-1* in regulating not only DSB formation, but also homology-mediated repair, we investigated the recombination frequency in different genetic intervals on chromosome I and V by monitoring SNP markers in Bristol/Hawaiian hybrids, which allowed us to assess both CO numbers and their position (Hillers and Villeneuve, 2009). We found a striking increase of COs in the central regions of both chromosomes (Fig. 6A-B), where COs are usually absent in the wild type (Lim et al., 2008). In addition, double and triple COs were observed, albeit at a low frequency. These results revealed that impaired *parg-1* function impacts the global levels and distribution of COs and weakens CO interference in *C. elegans*.

**Figure 6.**
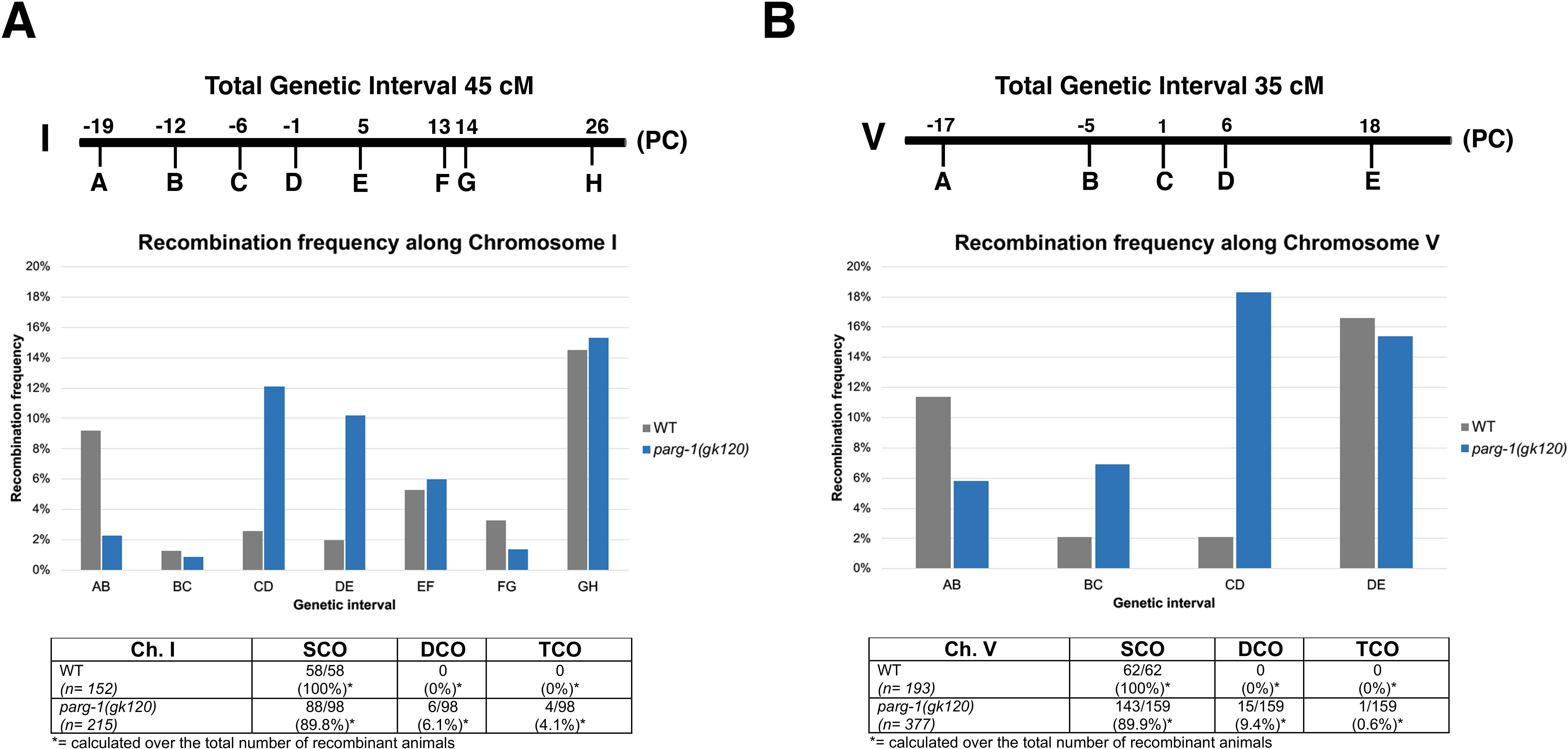
Loss of *parg-1* alters the recombination landscape and weakens CO interference. **(A)** Top: schematic representation of the genetic position of the SNPs employed to asses the recombination frequency on chromosome I. PC indicates the position of the pairing center. Middle: recombination frequencies assessed in each of the genetic intervals in wild type and *parg-1* null mutants. Bottom: table displaying number and percentage of single, double and triple crossovers (SCO, DCO and TCO respectively) in both genotypes. *n* indicates number of worms analyzed. **(B)** Same analysis as in A, performed for chromosome V.

### PARG-1 catalytic activity is dispensable for meiotic functions

We next sought to investigate whether PARG-1 catalytic activity is necessary to exert its function during meiosis. To this end, we generated a *parg-1* “catalytic-dead” mutant (referred to as *parg-1(cd)* hereafter) using CRISPR to mutate two glutamates in the catalytic domain (E554, 555A). These amino acids are conserved throughout evolution and were shown to be essential for PARG activity *in vitro* in both mammals and nematodes (Mortusewicz et al., 2011; Patel et al., 2005; St-Laurent et al., 2007). Immunostaining analysis in *parg-1(cd)* and *parg-1(cd)::GFP* revealed accumulation of PAR on meiotic chromosome axes as in *parg-1(gk120)* null mutants, indicating that also *in vivo* E554-E555 are necessary for PAR removal (Fig. S8). PARG-1^CD^::GFP was expressed and loaded in meiocytes (Fig. 7A) but displayed prolonged localization along the chromosomes in late pachytene cells, were PARG-1 normally is retained mostly at the short arm of the bivalent in control animals (Fig. 2). Western blot analysis showed that the overall levels of both PARG-1^CD^::GFP and untagged PARG-1^CD^ were indeed increased, ruling out possible artifacts due to the addition of GFP (Fig. 7B). The blots were also probed with anti-PAR antibodies and this confirmed that PAR accumulates in strains with compromised glycohydrolase activity (Fig. 7B).

**Figure 7.**
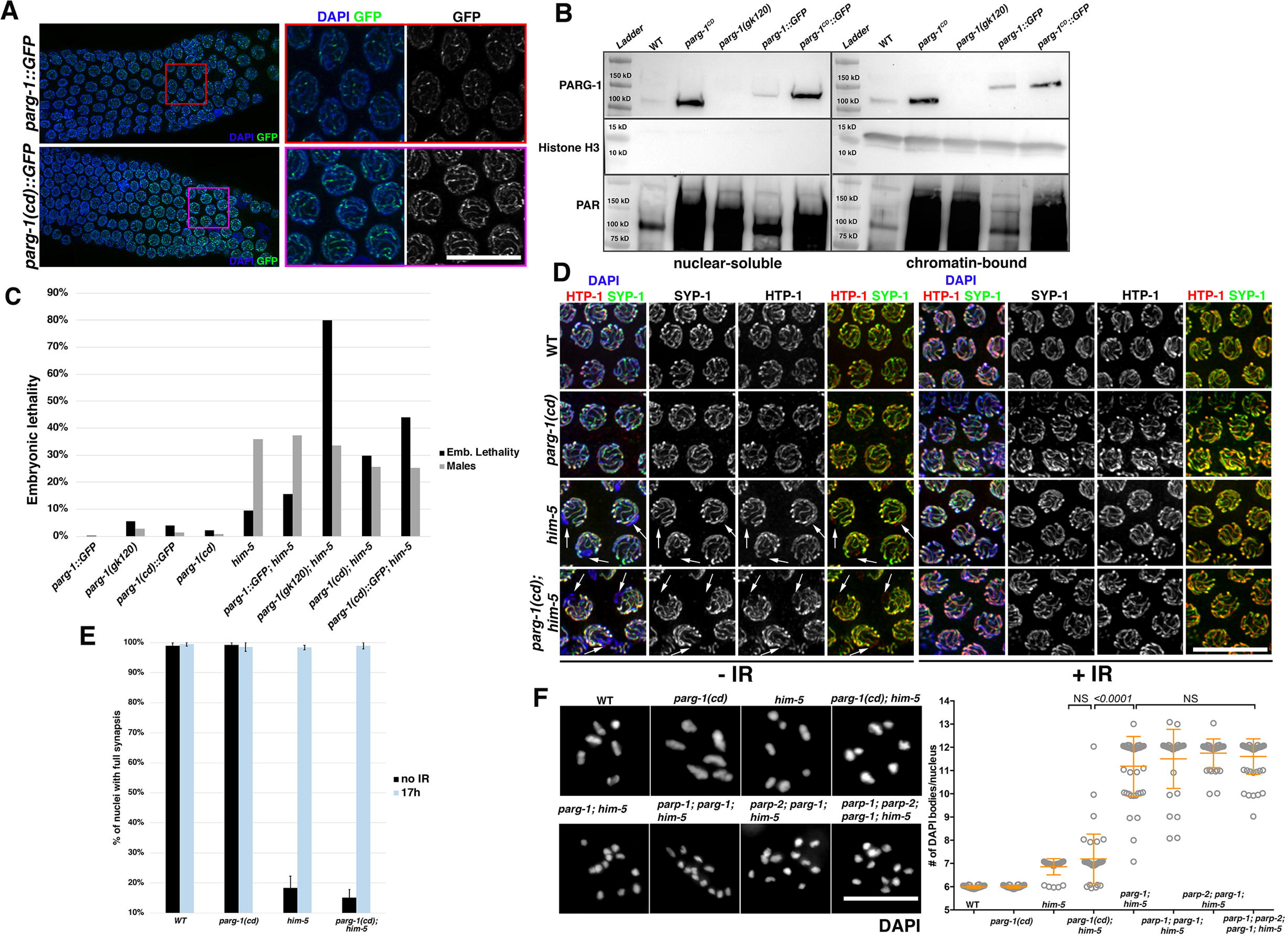
PARG-1 catalytic activity is dispensable for recombination. **(A)** Left: immunofluorescence of late pachytene nuclei showing elevated levels of PARG-1^CD^::GFP versus PARG-1::GFP and delayed redistribution. Right: insets depicting magnified nuclei from left. Scale bar 10 μm. **(B)** Western blots with fractionated extracts show higher PARG-1^CD^::GFP and PARG-1^CD^ abundance compared to controls. Anti-histone H3 was used as loading control for chromatin-bound fraction. Western blot (bottom) confirmed accumulation of PAR in both *parg-1(cd)* and *parg-1(gk120)* null mutants. **(C)** Blocking PARG-1 catalytic activity causes milder synergistic effects when combined with *him-5* mutants in contrast to *parg-1(gk120); him-5*. **(D)** De-synapsis of the X chromosome is fully rescued by IR exposure in *parg-1(cd); him-5* double mutants. Arrows indicate the unsynapsed X chromosome in the indicated genotypes before irradiation. Scale bar 10 μm. **(E)** Quantification of synapsis by SYP-1 and HTP-3 co-stainings in the indicated genotypes before and after IR exposure. Error bars indicated S.E.M. **(F)** DAPI-stainings (left) and quantification of DAPI-bodies in diakinesis nuclei (right) for the indicated genotypes. Error bars indicate standard deviation. Scale bar 5 μm.

To assess whether the catalytic activity of PARG-1 was required for the induction and/or repair of meiotic DSBs, we analyzed the *parg-1(cd); him-5* double mutants as described above. Offspring viability was only mildly reduced compared to *him-5* mutants (Fig. 7C). This indicates robust establishment of chiasmata in contrast to the *parg-1(gk120); him-5* double mutants. Similarly, X chromosome desynapsis (Fig. 7D-E) and its consequent nondisjunction (Fig. 7F) were rescued to the same extent upon IR exposure in both *parg-1(cd); him-5* and *him-5* mutants, unlike the lack of rescue in *parg-1(gk120); him-5* (Fig. 5A). These results suggest that PARG-1 loading onto chromosomes and/or a non-catalytic function of PARG-1 are essential to avert recombination defects in the absence of HIM-5. This interpretation was reinforced by the observation that the simultaneous removal of *parp-1* and *parp-2* did not rescue CO formation in *parg-1(gk120); him-5* mutants (Fig. 7F), indicating that CO defects are independent of PAR. Thus, we conclude that the glycohydrolase activity of PARG-1 is not required to promote induction of meiotic DSBs and their homologous recombination-mediated repair.

## Discussion

PARylation has been extensively studied in the context of the DNA damage response in mitotic mammalian cells, where it facilitates the repair of DNA lesions by promoting both the recruitment of repair factors and mediating local chromatin relaxation around damage sites (Gibson and Kraus, 2012; Gupte et al., 2017; Ray Chaudhuri and Nussenzweig, 2017; Weaver and Yang, 2013). In contrast to PARP1/2, the functions of PARG have been much less investigated due to the lack of a suitable model system, since PARG null mutants are embryonic lethal in mammals (O’Sullivan et al., 2019). We found that the *C. elegans* PARG-1 regulates DSB induction, in parallel to the so far known HIM-17/HIM-5/DSB-1/DSB-2-dependent routes. Moreover, our data demonstrate that PARG-1 regulates homology-directed repair of DSBs by operating within a functional module with HIM-5 to ensure the efficient conversion of recombination intermediates into post-recombination products, ultimately controlling global CO numbers.

Our cytological analysis, in combination with co-immunoprecipitation assays (Fig. 2), identified PARG-1 as an intrinsic component of the SC, where it is recruited *via* interaction with the chromosome axis protein HTP-3. Studies in mammalian mitotic cells reported nucleoplasmic localization of PARG and robust recruitment onto the DNA lesions induced by laser microirradiation (Kaufmann et al., 2017; Mortusewicz et al., 2011). The association with a meiosis-specific structure such as the SC therefore suggests distinct functional regulation in meiotic cells. Interestingly, PARG-1 retracts to the short arm of the bivalent and becomes enriched with SYP proteins at the presumptive CO sites in late pachytene nuclei (Fig. 2C-D), a localization also described for DNA repair and CO-promoting factors (Bhalla et al., 2008; Janisiw et al., 2018; Jantsch et al., 2004; Li et al., 2018). Nevertheless, abrogation of synapsis did not impair loading of PARG-1 along the chromosomes, a prerogative typically observed for axial rather than central components of the SC (de Carvalho et al., 2008; Goodyer et al., 2008; Martinez-Perez and Villeneuve, 2005; Zetka et al., 1999). This would suggest that PARG-1 may be targeted to both lateral and central elements of the SC or shift from the former to the latter upon CO-mediated chromosome remodeling. In support of a dynamic model of PARG-1 localization, PARG-1 was found in protein complexes both with HTP-1, HTP-3 and REC-8, all proteins localizing to chromosome axes (Goodyer et al., 2008; Martinez-Perez et al., 2008; Pasierbek et al., 2001), and also with SYP-3, which is a component of the central part of the SC (Smolikov et al., 2007). We believe that the localization of PARG-1 to the chromosome axes and its interaction with HTP-3 might hold crucial functional implications for promoting formation of meiotic DSBs. Many axial proteins, including *C. elegans* HTP-3, have been shown to directly influence abundance of DSBs during meiosis in several organisms (Goodyer et al., 2008; Kleckner, 2006), and therefore PARG-1 might exert its pro-DSB functions by operating from within the SC.

An activity of PARG-1 in promoting meiotic DSB formation by directly regulating pro-DSB factors is less likely, since the synergistic effects between *parg-1* and *him-17-him-5-dsb-2* mutants (Fig. 4, Fig. S5) clearly place PARG-1 in a parallel, distinct pathway. Consistently, expression and localization of PARG-1 and HIM-17, HIM-5 or DSB-2 were not mutually dependent (Fig. S5). We cannot rule out the possibility that PARG-1 may contribute to DSB formation by modulating SPO-11 activity or its recruitment to the presumptive DNA break sites. However, this is an unlikely scenario, since neither defects in bivalent formation nor RAD-51 loading were observed in *parg-1* mutants (Fig. 3; Fig. S2 and S3), as one might expect much more severe defects for as general loading problem. An additional argument in support of a model where interaction with HTP-3 might be key for PARG-1-mediated pro-DSB function, comes from its co-immunoprecipitation with MRE-11 (Fig. 4E), a proven interaction partner of HTP-3 (Goodyer et al., 2008). MRE-11 holds important roles in break resection and in *C. elegans* also break formation across species (Johzuka and Ogawa, 1995; Yin and Smolikove, 2013). MRE-11 has been invoked as a putative substrate intersected by HTP-3 function in inducing meiotic breaks (Goodyer et al., 2008). Therefore, PARG-1 might act together with HTP-3 and MRE-11 to ensure normal levels of breaks.

Our analysis also revealed that PARG-1, both independently and in combination with HIM-5, plays important roles in the global regulation of meiotic recombination. In fact, *parg-1* mutants show a profoundly perturbed recombination landscape, as distribution of COs displayed a marked shift towards the center of the autosomes (Fig. 6), a chromosome domain normally devoid of COs in wild-type animals (Barnes et al., 1995). This feature has also been observed in mutants with reduced levels bivalent formation or aberrant DSB repair (Jagut et al., 2016; Li et al., 2018; Meneely et al., 2012; Saito et al., 2013; Saito et al., 2009). Moreover, CO interference appeared weakened in absence of *parg-1*, suggesting a diminished stringency in the control of CO numbers.

The intermediates formed upon abrogation of *parg-1* function are nonetheless fully competent to be processed as COs, as long as HIM-5 function is preserved. In fact, while bivalent formation was fully restored in *parg-1; him-17* double mutants upon IR exposure (Fig. 4D) (arguing for a rescue of reduced DSB levels), diakinesis of irradiated *parg-1; him-5* mutant worms showed only a partial restoration of chiasmata, highlighting a repair defect as well (Fig. 4D). The mutual requirement of PARG-1 and HIM-5 in the reciprocal mutant background suggests the presence of a repair mechanism that relies on these two factors in order to efficiently complete interhomolog recombination repair. Both *him-5* and *dsb-2* exert regulatory functions on DNA repair pathway choice during gametogenesis (Macaisne et al., 2018) and our work also highlights *parg-1* as an important factor operating within such a process. This is also exemplified by our finding that diakinesis nuclei in *parg-1; mre-11(iow1)* display intact, well-shaped univalents both before and after IR exposure, in contrast to the chromatin clumps observed in the *mre-11(iow1)* separation of function mutant (Fig. 3) (Yin and Smolikove, 2013), indicating that PARG-1 can act as a switch in channeling DSB repair into multiple branches. Simultaneous abrogation of *parg-1* and *him-5* function caused much more severe aberrations than just reduced recombination: we found that IR exposure restored COSA-1 loading to the wildtype levels (six foci/nucleus) in pachytene cells (consistent with impaired break formation); nevertheless large portions of chromatin, possibly corresponding to whole chromosome pairs rather than local regions, were devoid of SYP-1/COSA-1 in many of these nuclei. These data indicate that additional COs have been designated on remaining, SC-associated chromosomes (Fig. 5).

Previous studies in *ex vivo* somatic cells suggested possible functions of PARG that are independent of its catalytic activity or PAR synthesis (Mortusewicz et al., 2011). Our data show that in catalytically impaired *parg-1(cd)* mutants, which consistently accumulate PAR as in *parg-1(gk120)* nulls (Fig. S8), the inactive protein was recruited at higher levels and displayed delayed redistribution along the chromosomes in late pachytene nuclei (Fig. 7A). This is in agreement with reports in mammalian cells showing that PARG^KD^ is recruited to laser-induced microirradiation sites with faster kinetics compared to PARG^WT^ and that this recruitment is only partially dependent on the PARP1 function (Mortusewicz et al., 2011). Strikingly, PARG-1^CD^::GFP was still capable of promoting chiasmata formation on the autosomes in *him-5* mutants: in fact, the embryonic viability and numbers of DAPI-bodies in *parg-1(cd); him-5* were comparable to *him-5* single mutants before and after IR exposure, and importantly, desynapsis was not observed. This suggests that the loading of PARG-1, rather than its enzymatic activity for PAR removal, was sufficient to induce DSBs and promote efficient bivalent formation in the presence of exogenous DSBs (Fig. 7). This was further corroborated by the fact that in the *parp-1; parp-2; parg-1(gk120); him-5* quadruple mutants, bivalent formation was not rescued, demonstrating that the roles exerted by PARG-1 in promoting DSB induction and meiotic repair are independent of PARylation.

Altogether, we demonstrate that the catalytic activity and the scaffolding properties of PARG are required for distinct cellular processes. Our study establishes a crucial role of PARG during meiotic prophase I in augmenting induction of meiotic DSBs and regulating their repair via HR in a metazoan model. Further studies are necessary to clarify whether PARG-1 recruitment affects the structure of the SC resulting in the modulation of DSB formation and recombination, or whether the presence of PARG-1 along the chromosomes influences the recruitment and dynamic behavior of other factors, which ultimately exert a regulatory role in DSB formation and recombination. Our work highlights the multifaceted aspects of PARG *in vivo* not simply as an enzyme mediating the catabolism of PAR, but also as a pivotal factor intersecting multiple functional branches acting during meiosis.

## Acknowledgements

We are grateful to E. Martinez-Perez, M. Zetka, A. Villeneuve, Y. Kim and S. Smolikove for strains and reagents; A. Graf for performing the microinjections; D. Slade for helpful comments and discussion throughout the development of this work; L. Krejčí, S. Uldrijan and D. Šmajs for sharing equipment. Some strains were provided by the CGC, which is funded by NIH Office of Research Infrastructure Programs (P40 OD010440). NS was funded by an Interdisciplinary Cancer Research (INDICAR) fellowship by the Mahlke-Obermann Stiftung and the European Union’s Seventh Framework Programme for Research, Technological Development under grant agreement no 609431 and a “Start-Up” grant from the Department of Biology of Masaryk University. VJ lab receives funding by the Austrian Science Fund (FWF; project no. P-31275-B28); AVH lab by DK Population Genetics (UW: W1225-B20), DK RNA (UW: W1207) and FWF URSPRUNG 2018 (I-1824-B22); JLY lab by NIGMS (2 R01 GM104007); MR was funded in part by an MWRI post-doctoral fellowship. We acknowledge the core facility CELLIM of CEITEC supported by the Czech-BioImaging large RI project (LM2018129 funded by MEYS CR) for their support with obtaining scientific data presented in this paper.

## Author contributions

NS designed the research and performed most of the experiments with the technical support of EJ; MR, FB and JLY generated some strains, analyzed diakinesis chromosomes, and performed the recombination assay on chromosome I; LPP and AVH analyzed whole genome sequencing data, which initiated the analysis of the catalytic-dead *parg-1* mutants; AB produced the *HA::rmh-1* tagged line; VJ provided logistic, infrastructure, resources and conceptual support; JLY, VJ and NS wrote the manuscript.

## Methods

### Genetics

Worms were cultured at 20°C according to standard conditions. The N_2_ strain was used as the wild-type control. We did not notice any significant differences between *him-17(e2707)* and *him-17(e2806)* alleles and the former has been employed for the majority of the experiments unless otherwise indicated. The *parp-1(ddr31)* is a full knock-out generated by CRISPR. A full list of the strains employed for this study are in Table S3.

### Screenings

L4 worms were individually plated and moved onto fresh plates every 12 hours for three days. Dead eggs were scored 24 hours after the mother had been removed and male progeny after three days. Embryonic lethality and male progeny were calculated as the fraction of unhatched eggs/total laid eggs and males/total hatched eggs respectively.

### Cytological procedures and image acquisition

For immunostaining experiments, synchronized worms of the indicated age were dissected and processed as previously described (Janisiw et al., 2018) except for detection of PARG-1::GFP and GFP::MSH-5. Briefly, worms were dissected in PBS and immediately placed in liquid nitrogen. Slides were placed in cold methanol at −20°C for 1 and then fixed with 2% PFA in 0.1M K_2_HPO_4_ (pH 7.4) for 10 in a humid chamber at room temperature. Samples were then processed as for regular staining. For GFP::MSH-5 detection, worms were dissected and fixed in 2.5% PFA for 2 at room temperature and then freeze-cracked in liquid nitrogen. Slides were placed in absolute ethanol at −20°C for 10 and then washed in 1x PBST. DAPI staining was performed as for normal staining and GFP was directly acquired without employing a primary anti-GFP antibody. For quantification of PAR (Figure 1E), samples were acquired with identical settings and equally adjusted in Fiji. Gonads were divided into seven equal regions from mitotic tip to diplotene entry and a circle of fixed area was employed to assess absolute fluorescence in each nucleus with Fiji as in (Janisiw et al., 2018). Between two and three germlines for each genotype were used for quantification. Number of nuclei scored was (from zone 1 to 7): WT (97, 129, 115, 107, 102, 74, 45), *parg-1(gk120)* (129, 136, 140, 129, 130, 96, 82), *parp-1(ddr31); parg-1(gk120)* (93, 113, 123, 129, 102, 69, 44), *parp-2(ok344); parg-1* (190, 263, 215, 179, 179, 125, 72), *parp-1(ddr31); parp-2(ok344); parg-1(gk120)* (107, 140, 134, 144, 126, 103, 76).

For quantification of RAD-51 foci, gonads were divided into seven equal regions from the mitotic tip to the diplotene entry and number of RAD-51 foci was counted in each nucleus. Number of nuclei analyzed is reported in Table S2.

Quantification of phospho-SUN-1^S8^ extension was performed as in (Link et al., 2018). Most images were captured using a Delta Vision system equipped with an Olympus IX-71 microscope and a Roper CoolSNAP HQ2 camera with Z-stack set at 0.25 μm of thickness. Images in Figure 7D and 7F, were acquired with a Delta Vision system equipped with an Evolve 512 EMCCD Camera and an upright fluorescence microscope Zeiss AxioImager.Z2 equipped with a Hamamatsu ORCA Flash 4.0, sCMOS sensor camera respectively, using UPlanSApo 100x/1.4 Oil objective. All images were deconvolved using Softworx (Applied Precision) except for images in Figure 7F, which are non-deconvolved.

### Antibodies

The following antibodies at the indicated dilutions were employed for immunolocalization studies: rabbit polyclonal anti HA (SIGMA, 1:1000), rabbit polyclonal anti OLLAS (Genscript, 1:1500), rabbit polyclonal anti PAR (Trevigen, 1:1000), mouse monoclonal anti GFP (Roche, 1:500), guinea pig polyclonal anti HTP-3 (1:500) (Goodyer et al., 2008), guinea pig polyclonal anti HTP-3 (1:750) (Y. Kim lab), chicken polyclonal anti SYP-1 (1:500) (Silva et al., 2014), rabbit polyclonal anti HTP-1 (1:500) (Martinez-Perez et al., 2008), rabbit polyclonal anti RAD-51 (Novus, 1:10,000), guinea pig polyclonal anti phospho-SUN-1^S8^ (1:750) (Woglar et al., 2013), rabbit polyclonal anti DSB-2 (1:5000) (Rosu et al., 2013), guinea pig polyclonal anti XND-1 (1:2500) (Wagner et al., 2010), mouse monoclonal anti H3K4me2 (Millipore, 1:250). All the secondary antibodies were Alexafluor-conjugated and used at 1:500. The following antibodies at the indicated dilutions were employed in western blot analysis: rabbit polyclonal anti HA (SIGMA, 1:3000), mouse monoclonal anti HA (Cell Signalling, 1:1000), mouse monoclonal anti PARG-1 (this study, 1:500), chicken polyclonal anti GFP (Abcam, 1:4000), rabbit polyclonal anti Histone H3 (Abcam, 1:100,000), goat polyclonal anti actin (Santa Cruz, 1:3000), mouse monoclonal anti Tubulin (Thermofisher, 1:2000), mouse monoclonal anti GAPDH (Ambion, 1:5000). Secondary antibodies were purchased from Thermofisher and were HRP-conjugated.

### Biochemistry

Fractionated protein extracts were produced as previously described (Silva et al., 2014) and co-immunoprecipitation assays and Western Blot were performed as previously shown (Janisiw et al., 2018). At least 500μg of nuclear extract (pooled nuclear-soluble and chromatin-bound fractions) were used for IPs. Agarose GFP-traps (Chromotek) were employed for pull downs following manufacturer instructions. Buffer D (20mM HEPES pH 7.9, 150mM KCl, 20% glycerol, 0.2mM EDTA, 0.2% Triton X-100 and complete protease inhibitor) was used for incubation with beads and washes.

### Generation of anti-PARG-1 antibody

To generate the mouse monoclonal anti PARG-1(2D4) antibody, the cDNA encoding for residues 1-350 of *C. elegans* PARG-1 (isoform A) was generated by gene synthesis (IDT) and then cloned into pCoofy31 in frame with a C-ter 6xHis tail. The resulting plasmid was expressed in *E. coli* BL21 cells according to standard procedures and 1 mg of purified protein was used to immunize three mice in the “in-house” monoclonal antibody facility at Max Perutz Laboratories (https://www.maxperutzlabs.ac.at/research/facilities/monoclonal-antibody-facility). Raw sera were screened by western blot employing extracts produced from WT, *parg-1(gk120)* and *parg-1::GFP* worms in order to identify immunoreactive bands against PARG-1. Spleen cells from one mouse were fused with myeloma cells to generate hybridoma cell lines and mixed clones were successively diluted to gain monoclonal line 2D4, from which the antibody was harvested. Antibody specificity was assessed by western blot, where an immune reactive band of the expected Mw of approximately 90kDa in WT but not in *parg-1* mutant worms was detected (Fig.2A).

### Irradiation

Irradiation assays were performed as previously described (Janisiw et al., 2018). For quantification of synapsis and OLLAS::COSA-1 foci number in late pachytene nuclei, worms were dissected at the indicated time after irradiation and quantification was performed in the last seven rows of nuclei before diplotene entry. For quantification of HA::RMH-1 and GFP::MSH-5 in Figure 4, worms were dissected 8h post-IR and gonads from transition zone to late pachytene were divided into five equal regions and number of foci/nucleus was assessed. For diakinesis analysis, worms were dissected 24-27 hours post irradiation. The dose employed for all irradiation experiments was 10 Gy. Number of nuclei analyzed for each condition are reported in Table S1 and S2.

### CRISPR-Cas9 genome editing

Generation of tagged or mutated lines was performed as previously described (Janisiw et al., 2018). Briefly, to tag endogenous locus of *parg-1*, GFP was amplified by PCR with primers carrying 25 base pairs of homology to the left and right side of the STOP codon of *parg-1* gene. To generate the PARG-1^E555,556A^ catalytic dead mutant, a synthetic ultramer (IDT) was employed, in which we included silent mutations to produce an Alu I restriction site for screening purposes. The mutations were generated in both WT and *parg-1::GFP* genetic backgrounds. To elicit a full knock-out of *parp-1*, we employed two sgRNAs targeting the beginning and the end of the gene. The *him-17::3xHA* and *him-5::3xHA* were generated by employing synthetic DNA ultramers (IDT) and N2 worms were injected. All the tagged lines carried a 5x-Gly linker between the tag and the coding region. The *parg-1^gk120^* line carries the same deletion present in the VC130 strain, which we generated in both WT and CB4856 strains. All the lines generated by CRISPR were outcrossed to WT worms at least twice before use.

### Recombination Assay

The recombination landscape was assessed following the same strategy as in (Hillers and Villeneuve, 2003), by exploiting different Dra I digestion pattern of SNPs present in the Bristol and Hawaiian genetic backgrounds. Briefly, *parg-1(gk120)* and *parg-1(cd)* mutations were generated by CRISPR in both the N2 (Bristol) and CB4856 (Hawaii) strains. Bristol/Hawaiian F1 hermaphrodite hybrids carrying the indicated mutations were backcrossed to Bristol males carrying a tdTomato fluorescent reporter expressed in the soma in order to monitor recombination frequency in the oocytes. The relevant regions containing the SNPs for chromosomes I and V in the indicated genetic intervals were amplified by PCR and the products digested with Dra I to monitor recombination patterns. Data presented in Figure 6 refer to the total number of worms analyzed in independent replicates.

## Supplementary Figure Legends

**Figure S1. PAR accumulates in absence of endogenous DSBs.**

Immunostaining analysis showing accumulation of PAR in *parg-1* mutants even in absence of SPO-11-induced DSBs. *parg-1::GFP* does not accumulate PAR, indicating functionality of the fusion protein. Scale bar 10 μm.

**Figure S2. PARG-1 is dispensable for synapsis and CO formation.**

**(A)** Staining of SYP-1 and HTP-3 shows no abnormalities in *parg-1(gk120)* mutants compared to wild-type (WT) animals. **(B)** Quantification of DAPI-bodies (left) and representative examples (right) of diakinesis nuclei in the indicated genotypes. Scale bar 2 μm.

**Figure S3. Abrogation of parg-1 function causes reduced amounts of RAD-51 foci in *him*-5 and *him*-17 mutants.**

**(A)** Quantification of RAD-51 foci throughout the germline in the indicated genotypes (top) and representative images of mid-pachytene nuclei stained with DAPI and anti RAD-51 (bottom). Error bars indicate S.E.M. Statistical analysis was performed using non-parametric two-tailed Mann-Whitney test. (\*\*\**p<0.0001*; \*\**p= 0.0036*). **(B)** Removal of *parg-1* causes synthetic lethality in DSB-defective mutants. **(C)** The CO defects in *dsb-2* mutants are exacerbated by lack of PARG-1.

**Figure S4. Loss of PARG-1 impairs loading of pro-CO factors in *him-5* and *him-17* mutants.**

**(A)** Late pachytene nuclei of different genotypes stained for GFP (MSH-5) before and after IR. **(B)** Late pachytene nuclei of different genotypes stained for HA (RMH-1) and OLLAS (COSA-1) before and after IR Scale bar 10 μm.

**Figure S5. PARG-1 and pro-DSB factors display a mutually independent loading.**

**(A)** Viability and male progeny assessment revealed full functionality of *him-5::3xHA* and *him-17::3xHA* tagged lines compared to respective mutant backgrounds. **(B)** HIM-5 shows normal loading in *parg-1* mutant germlines. Scale bar 20 μm. **(C)** HIM-17 and XND-1 do not display loading abnormalities in *parg-1* mutants. Scale bar 20 μm. **(D)** *parg-1* is not required for DSB-2 loading. Scale bar 20 μm. **(E)** Loading of PARG-1 is not dependent on *him-5*, *him-17* and *dsb-2*. Scale bar 20 μm.

**Figure S6. Loss of *parg-2* does not cause synthetic phenotypes with *him-5*.**

**(A)** Quantification of DAPI-bodies in diakinesis nuclei of the indicated genotypes before and after IR exposure. **(B)** Representative images of diakinesis nuclei of the indicated genotypes stained with DAPI. Scale bar 5 μm.

**Figure S7. IR does not fully suppress accumulation of pSUN-1^S8^ in *parg-1; him-5* double mutants.**

**(A)** Quantification of pSUN-1^S8^-positive nuclei cell rows in the indicated genotypes before and after IR exposure. **(B)** Representative whole-mount gonads of the indicated genotypes before and after irradiation, stained for pSUN-1^S8^ and DAPI. Scale bar 30 μm.

**Figure S8. PARG-1^E554,555A^ is a “catalytic-dead” mutant of PARG-1.**

Left: whole-mount germlines of the indicated genotypes stained with anti-PAR antibodies and DAPI. Right: PAR staining. Scale bar 20 μm.

## References

Alpi, A., Pasierbek, P., Gartner, A., and Loidl, J. (2003). Genetic and cytological characterization of the recombination protein RAD-51 in Caenorhabditis elegans. Chromosoma 112, 6–16.

Ame, J.C., Fouquerel, E., Gauthier, L.R., Biard, D., Boussin, F.D., Dantzer, F., de Murcia, G., and Schreiber, V. (2009). Radiation-induced mitotic catastrophe in PARG-deficient cells. J Cell Sci 122, 1990–2002.

Bae, W., Park, J.H., Lee, M.H., Park, H.W., and Koo, H.S. (2019). Hypersensitivity to DNA double-strand breaks associated with PARG deficiency is suppressed by exo-1 and polq-1 mutations in Caenorhabditis elegans. FEBS J.

Barnes, T.M., Kohara, Y., Coulson, A., and Hekimi, S. (1995). Meiotic recombination, noncoding DNA and genomic organization in Caenorhabditis elegans. Genetics 141, 159–179.

Bhalla, N., Wynne, D.J., Jantsch, V., and Dernburg, A.F. (2008). ZHP-3 acts at crossovers to couple meiotic recombination with synaptonemal complex disassembly and bivalent formation in C. elegans. PLoS Genet 4, e1000235.

Byrne, A.B., McWhirter, R.D., Sekine, Y., Strittmatter, S.M., Miller, D.M., and Hammarlund, M. (2016). Inhibiting poly(ADP-ribosylation) improves axon regeneration. Elife 5.

Cao, L., Alani, E., and Kleckner, N. (1990). A pathway for generation and processing of double-strand breaks during meiotic recombination in S. cerevisiae. Cell 61, 1089–1101.

Chin, G.M., and Villeneuve, A.M. (2001). C. elegans mre-11 is required for meiotic recombination and DNA repair but is dispensable for the meiotic G(2) DNA damage checkpoint. Genes Dev 15, 522–534.

Colaiacovo, M.P., MacQueen, A.J., Martinez-Perez, E., McDonald, K., Adamo, A., La Volpe, A., and Villeneuve, A.M. (2003). Synaptonemal complex assembly in C. elegans is dispensable for loading strand-exchange proteins but critical for proper completion of recombination. Dev Cell 5, 463–474.

Crawley, O., Barroso, C., Testori, S., Ferrandiz, N., Silva, N., Castellano-Pozo, M., Jaso-Tamame, A.L., and Martinez-Perez, E. (2016). Cohesin-interacting protein WAPL-1 regulates meiotic chromosome structure and cohesion by antagonizing specific cohesin complexes. Elife 5, e10851.

Dantzer, F., Mark, M., Quenet, D., Scherthan, H., Huber, A., Liebe, B., Monaco, L., Chicheportiche, A., Sassone-Corsi, P., de Murcia, G., et al. (2006). Poly(ADP-ribose) polymerase-2 contributes to the fidelity of male meiosis I and spermiogenesis. Proc Natl Acad Sci U S A 103, 14854–14859.

de Carvalho, C.E., Zaaijer, S., Smolikov, S., Gu, Y., Schumacher, J.M., and Colaiacovo, M.P. (2008). LAB-1 antagonizes the Aurora B kinase in C. elegans. Genes Dev 22, 2869–2885.

de los Santos, T., Hunter, N., Lee, C., Larkin, B., Loidl, J., and Hollingsworth, N.M. (2003). The Mus81/Mms4 endonuclease acts independently of double-Holliday junction resolution to promote a distinct subset of crossovers during meiosis in budding yeast. Genetics 164, 81–94.

Dequen, F., Gagnon, S.N., and Desnoyers, S. (2005). Ionizing radiations in Caenorhabditis elegans induce poly(ADP-ribosyl)ation, a conserved DNA-damage response essential for survival. DNA Repair (Amst) 4, 814–825.

Gagnon, S.N., Hengartner, M.O., and Desnoyers, S. (2002). The genes pme-1 and pme-2 encode two poly(ADP-ribose) polymerases in Caenorhabditis elegans. Biochem J 368, 263–271.

Gibson, B.A., and Kraus, W.L. (2012). New insights into the molecular and cellular functions of poly(ADP-ribose) and PARPs. Nat Rev Mol Cell Biol 13, 411–424.

Goodyer, W., Kaitna, S., Couteau, F., Ward, J.D., Boulton, S.J., and Zetka, M. (2008). HTP-3 links DSB formation with homolog pairing and crossing over during C. elegans meiosis. Dev Cell 14, 263–274.

Gupte, R., Liu, Z., and Kraus, W.L. (2017). PARPs and ADP-ribosylation: recent advances linking molecular functions to biological outcomes. Genes Dev 31, 101–126.

Hayashi, M., Chin, G.M., and Villeneuve, A.M. (2007). C. elegans germ cells switch between distinct modes of double-strand break repair during meiotic prophase progression. PLoS Genet 3, e191.

Hillers, K.J., and Villeneuve, A.M. (2003). Chromosome-wide control of meiotic crossing over in C. elegans. Curr Biol 13, 1641–1647.

Hillers, K.J., and Villeneuve, A.M. (2009). Analysis of meiotic recombination in Caenorhabditis elegans. Methods Mol Biol 557, 77–97.

Hodgkin, J., Horvitz, H.R., and Brenner, S. (1979). Nondisjunction Mutants of the Nematode CAENORHABDITIS ELEGANS. Genetics 91, 67–94.

Jagut, M., Hamminger, P., Woglar, A., Millonigg, S., Paulin, L., Mikl, M., Dello Stritto, M.R., Tang, L., Habacher, C., Tam, A., et al. (2016). Separable Roles for a Caenorhabditis elegans RMI1 Homolog in Promoting and Antagonizing Meiotic Crossovers Ensure Faithful Chromosome Inheritance. PLoS Biol 14, e1002412.

Janisiw, E., Dello Stritto, M.R., Jantsch, V., and Silva, N. (2018). BRCA1-BARD1 associate with the synaptonemal complex and pro-crossover factors and influence RAD-51 dynamics during Caenorhabditis elegans meiosis. PLoS Genet 14, e1007653.

Jantsch, V., Pasierbek, P., Mueller, M.M., Schweizer, D., Jantsch, M., and Loidl, J. (2004). Targeted gene knockout reveals a role in meiotic recombination for ZHP-3, a Zip3-related protein in Caenorhabditis elegans. Mol Cell Biol 24, 7998–8006.

Johzuka, K., and Ogawa, H. (1995). Interaction of Mre11 and Rad50: two proteins required for DNA repair and meiosis-specific double-strand break formation in Saccharomyces cerevisiae. Genetics 139, 1521–1532.

Kaufmann, T., Grishkovskaya, I., Polyansky, A.A., Kostrhon, S., Kukolj, E., Olek, K.M., Herbert, S., Beltzung, E., Mechtler, K., Peterbauer, T., et al. (2017). A novel non-canonical PIP-box mediates PARG interaction with PCNA. Nucleic Acids Res 45, 9741–9759.

Keeney, S., Giroux, C.N., and Kleckner, N. (1997). Meiosis-specific DNA double-strand breaks are catalyzed by Spo11, a member of a widely conserved protein family. Cell 88, 375–384.

Kelly, K.O., Dernburg, A.F., Stanfield, G.M., and Villeneuve, A.M. (2000). Caenorhabditis elegans msh-5 is required for both normal and radiation-induced meiotic crossing over but not for completion of meiosis. Genetics 156, 617–630.

Kleckner, N. (2006). Chiasma formation: chromatin/axis interplay and the role(s) of the synaptonemal complex. Chromosoma 115, 175–194.

Koh, D.W., Lawler, A.M., Poitras, M.F., Sasaki, M., Wattler, S., Nehls, M.C., Stoger, T., Poirier, G.G., Dawson, V.L., and Dawson, T.M. (2004). Failure to degrade poly(ADP-ribose) causes increased sensitivity to cytotoxicity and early embryonic lethality. Proc Natl Acad Sci U S A 101, 17699–17704.

Li, Q., Saito, T.T., Martinez-Garcia, M., Deshong, A.J., Nadarajan, S., Lawrence, K.S., Checchi, P.M., Colaiacovo, M.P., and Engebrecht, J. (2018). The tumor suppressor BRCA1-BARD1 complex localizes to the synaptonemal complex and regulates recombination under meiotic dysfunction in Caenorhabditis elegans. PLoS Genet 14, e1007701.

Libuda, D.E., Uzawa, S., Meyer, B.J., and Villeneuve, A.M. (2013). Meiotic chromosome structures constrain and respond to designation of crossover sites. Nature 502, 703–706.

Lim, J.G., Stine, R.R., and Yanowitz, J.L. (2008). Domain-specific regulation of recombination in Caenorhabditis elegans in response to temperature, age and sex. Genetics 180, 715–726.

Link, J., Paouneskou, D., Velkova, M., Daryabeigi, A., Laos, T., Labella, S., Barroso, C., Pacheco Pinol, S., Montoya, A., Kramer, H., et al. (2018). Transient and Partial Nuclear Lamina Disruption Promotes Chromosome Movement in Early Meiotic Prophase. Dev Cell 45, 212–225 e217.

Macaisne, N., Kessler, Z., and Yanowitz, J.L. (2018). Meiotic Double-Strand Break Proteins Influence Repair Pathway Utilization. Genetics 210, 843–856.

Machovina, T.S., Mainpal, R., Daryabeigi, A., McGovern, O., Paouneskou, D., Labella, S., Zetka, M., Jantsch, V., and Yanowitz, J.L. (2016). A Surveillance System Ensures Crossover Formation in C. elegans. Curr Biol 26, 2873–2884.

MacQueen, A.J., Colaiacovo, M.P., McDonald, K., and Villeneuve, A.M. (2002). Synapsis-dependent and -independent mechanisms stabilize homolog pairing during meiotic prophase in C. elegans. Genes Dev 16, 2428–2442.

Martinez-Perez, E., Schvarzstein, M., Barroso, C., Lightfoot, J., Dernburg, A.F., and Villeneuve, A.M. (2008). Crossovers trigger a remodeling of meiotic chromosome axis composition that is linked to two-step loss of sister chromatid cohesion. Genes Dev 22, 2886–2901.

Martinez-Perez, E., and Villeneuve, A.M. (2005). HTP-1-dependent constraints coordinate homolog pairing and synapsis and promote chiasma formation during C. elegans meiosis. Genes Dev 19, 2727–2743.

Mateo, A.R., Kessler, Z., Jolliffe, A.K., McGovern, O., Yu, B., Nicolucci, A., Yanowitz, J.L., and Derry, W.B. (2016). The p53-like Protein CEP-1 Is Required for Meiotic Fidelity in C. elegans. Curr Biol 26, 1148–1158.

Meneely, P.M., McGovern, O.L., Heinis, F.I., and Yanowitz, J.L. (2012). Crossover distribution and frequency are regulated by him-5 in Caenorhabditis elegans. Genetics 190, 1251–1266.

Menissier de Murcia, J., Ricoul, M., Tartier, L., Niedergang, C., Huber, A., Dantzer, F., Schreiber, V., Ame, J.C., Dierich, A., LeMeur, M., et al. (2003). Functional interaction between PARP-1 and PARP-2 in chromosome stability and embryonic development in mouse. EMBO J 22, 2255–2263.

Meyer-Ficca, M.L., Meyer, R.G., Coyle, D.L., Jacobson, E.L., and Jacobson, M.K. (2004). Human poly(ADP-ribose) glycohydrolase is expressed in alternative splice variants yielding isoforms that localize to different cell compartments. Exp Cell Res 297, 521–532.

Mortusewicz, O., Fouquerel, E., Ame, J.C., Leonhardt, H., and Schreiber, V. (2011). PARG is recruited to DNA damage sites through poly(ADP-ribose)- and PCNA-dependent mechanisms. Nucleic Acids Res 39, 5045–5056.

O’Sullivan, J., Tedim Ferreira, M., Gagne, J.P., Sharma, A.K., Hendzel, M.J., Masson, J.Y., and Poirier, G.G. (2019). Emerging roles of eraser enzymes in the dynamic control of protein ADP-ribosylation. Nat Commun 10, 1182.

Ohashi, S., Kanai, M., Hanai, S., Uchiumi, F., Maruta, H., Tanuma, S., and Miwa, M. (2003). Subcellular localization of poly(ADP-ribose) glycohydrolase in mammalian cells. Biochem Biophys Res Commun 307, 915–921.

Paix, A., Folkmann, A., Rasoloson, D., and Seydoux, G. (2015). High Efficiency, Homology-Directed Genome Editing in Caenorhabditis elegans Using CRISPR-Cas9 Ribonucleoprotein Complexes. Genetics 201, 47-54.

Pasierbek, P., Jantsch, M., Melcher, M., Schleiffer, A., Schweizer, D., and Loidl, J. (2001). A Caenorhabditis elegans cohesion protein with functions in meiotic chromosome pairing and disjunction. Genes Dev 15, 1349–1360.

Patel, C.N., Koh, D.W., Jacobson, M.K., and Oliveira, M.A. (2005). Identification of three critical acidic residues of poly(ADP-ribose) glycohydrolase involved in catalysis: determining the PARG catalytic domain. Biochem J 388, 493–500.

Pattabiraman, D., Roelens, B., Woglar, A., and Villeneuve, A.M. (2017). Meiotic recombination modulates the structure and dynamics of the synaptonemal complex during C. elegans meiosis. PLoS Genet 13, e1006670.

Penkner, A., Portik-Dobos, Z., Tang, L., Schnabel, R., Novatchkova, M., Jantsch, V., and Loidl, J. (2007). A conserved function for a Caenorhabditis elegans Com1/Sae2/CtIP protein homolog in meiotic recombination. EMBO J 26, 5071–5082.

Penkner, A.M., Fridkin, A., Gloggnitzer, J., Baudrimont, A., Machacek, T., Woglar, A., Csaszar, E., Pasierbek, P., Ammerer, G., Gruenbaum, Y., et al. (2009). Meiotic chromosome homology search involves modifications of the nuclear envelope protein Matefin/SUN-1. Cell 139, 920–933.

Ray Chaudhuri, A., and Nussenzweig, A. (2017). The multifaceted roles of PARP1 in DNA repair and chromatin remodelling. Nat Rev Mol Cell Biol 18, 610–621.

Reddy, K.C., and Villeneuve, A.M. (2004). C. elegans HIM-17 links chromatin modification and competence for initiation of meiotic recombination. Cell 118, 439–452.

Reichman, R., Shi, Z., Malone, R., and Smolikove, S. (2018). Mitotic and Meiotic Functions for the SUMOylation Pathway in the Caenorhabditis elegans Germline. Genetics 208, 1421–1441.

Reuben, M., and Lin, R. (2002). Germline X chromosomes exhibit contrasting patterns of histone H3 methylation in Caenorhabditis elegans. Dev Biol 245, 71–82.

Rog, O., and Dernburg, A.F. (2015). Direct Visualization Reveals Kinetics of Meiotic Chromosome Synapsis. Cell Rep.

Rosu, S., Libuda, D.E., and Villeneuve, A.M. (2011). Robust crossover assurance and regulated interhomolog access maintain meiotic crossover number. Science 334, 1286–1289.

Rosu, S., Zawadzki, K.A., Stamper, E.L., Libuda, D.E., Reese, A.L., Dernburg, A.F., and Villeneuve, A.M. (2013). The C. elegans DSB-2 protein reveals a regulatory network that controls competence for meiotic DSB formation and promotes crossover assurance. PLoS Genet 9, e1003674.

Saito, T.T., Lui, D.Y., Kim, H.M., Meyer, K., and Colaiacovo, M.P. (2013). Interplay between structure-specific endonucleases for crossover control during Caenorhabditis elegans meiosis. PLoS Genet 9, e1003586.

Saito, T.T., Youds, J.L., Boulton, S.J., and Colaiacovo, M.P. (2009). Caenorhabditis elegans HIM-18/SLX-4 interacts with SLX-1 and XPF-1 and maintains genomic integrity in the germline by processing recombination intermediates. PLoS Genet 5, e1000735.

Serrentino, M.E., and Borde, V. (2012). The spatial regulation of meiotic recombination hotspots: are all DSB hotspots crossover hotspots? Exp Cell Res 318, 1347–1352.

Silva, N., Ferrandiz, N., Barroso, C., Tognetti, S., Lightfoot, J., Telecan, O., Encheva, V., Faull, P., Hanni, S., Furger, A., et al. (2014). The fidelity of synaptonemal complex assembly is regulated by a signaling mechanism that controls early meiotic progression. Dev Cell 31, 503–511.

Slade, D. (2019). Mitotic functions of poly(ADP-ribose) polymerases. Biochem Pharmacol.

Smolikov, S., Eizinger, A., Schild-Prufert, K., Hurlburt, A., McDonald, K., Engebrecht, J., Villeneuve, A.M., and Colaiacovo, M.P. (2007). SYP-3 restricts synaptonemal complex assembly to bridge paired chromosome axes during meiosis in Caenorhabditis elegans. Genetics 176, 2015–2025.

St-Laurent, J.F., Gagnon, S.N., Dequen, F., Hardy, I., and Desnoyers, S. (2007). Altered DNA damage response in Caenorhabditis elegans with impaired poly(ADP-ribose) glycohydrolases genes expression. DNA Repair (Amst) 6, 329–343.

Stamper, E.L., Rodenbusch, S.E., Rosu, S., Ahringer, J., Villeneuve, A.M., and Dernburg, A.F. (2013). Identification of DSB-1, a protein required for initiation of meiotic recombination in Caenorhabditis elegans, illuminates a crossover assurance checkpoint. PLoS Genet 9, e1003679.

Sun, H., Treco, D., Schultes, N.P., and Szostak, J.W. (1989). Double-strand breaks at an initiation site for meiotic gene conversion. Nature 338, 87–90.

Tsai, C.J., Mets, D.G., Albrecht, M.R., Nix, P., Chan, A., and Meyer, B.J. (2008). Meiotic crossover number and distribution are regulated by a dosage compensation protein that resembles a condensin subunit. Genes Dev 22, 194–211.

Wagner, C.R., Kuervers, L., Baillie, D.L., and Yanowitz, J.L. (2010). xnd-1 regulates the global recombination landscape in Caenorhabditis elegans. Nature 467, 839–843.

Weaver, A.N., and Yang, E.S. (2013). Beyond DNA Repair: Additional Functions of PARP-1 in Cancer. Front Oncol 3, 290.

Winstall, E., Affar, E.B., Shah, R., Bourassa, S., Scovassi, I.A., and Poirier, G.G. (1999). Preferential perinuclear localization of poly(ADP-ribose) glycohydrolase. Exp Cell Res 251, 372–378.

Woglar, A., Daryabeigi, A., Adamo, A., Habacher, C., Machacek, T., La Volpe, A., and Jantsch, V. (2013). Matefin/SUN-1 phosphorylation is part of a surveillance mechanism to coordinate chromosome synapsis and recombination with meiotic progression and chromosome movement. PLoS Genet 9, e1003335.

Yin, Y., and Smolikove, S. (2013). Impaired resection of meiotic double-strand breaks channels repair to nonhomologous end joining in Caenorhabditis elegans. Mol Cell Biol 33, 2732–2747.

Yokoo, R., Zawadzki, K.A., Nabeshima, K., Drake, M., Arur, S., and Villeneuve, A.M. (2012). COSA-1 reveals robust homeostasis and separable licensing and reinforcement steps governing meiotic crossovers. Cell 149, 75–87.

Youds, J.L., Mets, D.G., McIlwraith, M.J., Martin, J.S., Ward, J.D., NJ, O.N., Rose, A.M., West, S.C., Meyer, B.J., and Boulton, S.J. (2010). RTEL-1 enforces meiotic crossover interference and homeostasis. Science 327, 1254–1258.

Zalevsky, J., MacQueen, A.J., Duffy, J.B., Kemphues, K.J., and Villeneuve, A.M. (1999). Crossing over during Caenorhabditis elegans meiosis requires a conserved MutS-based pathway that is partially dispensable in budding yeast. Genetics 153, 1271–1283.

Zetka, M.C., Kawasaki, I., Strome, S., and Muller, F. (1999). Synapsis and chiasma formation in Caenorhabditis elegans require HIM-3, a meiotic chromosome core component that functions in chromosome segregation. Genes Dev 13, 2258–2270.

Zickler, D., and Kleckner, N. (1999). Meiotic chromosomes: integrating structure and function. Annu Rev Genet 33, 603–754.

Zickler, D., and Kleckner, N. (2015). Recombination, Pairing, and Synapsis of Homologs during Meiosis. Cold Spring Harb Perspect Biol 7.

