## Supplementary figures and images for "Poly(ADP-ribose) glycohydrolase promotes formation and homology-directed repair of meiotic DNA double-strand breaks independent of its catalytic activity"

### Supplemental Fig 1

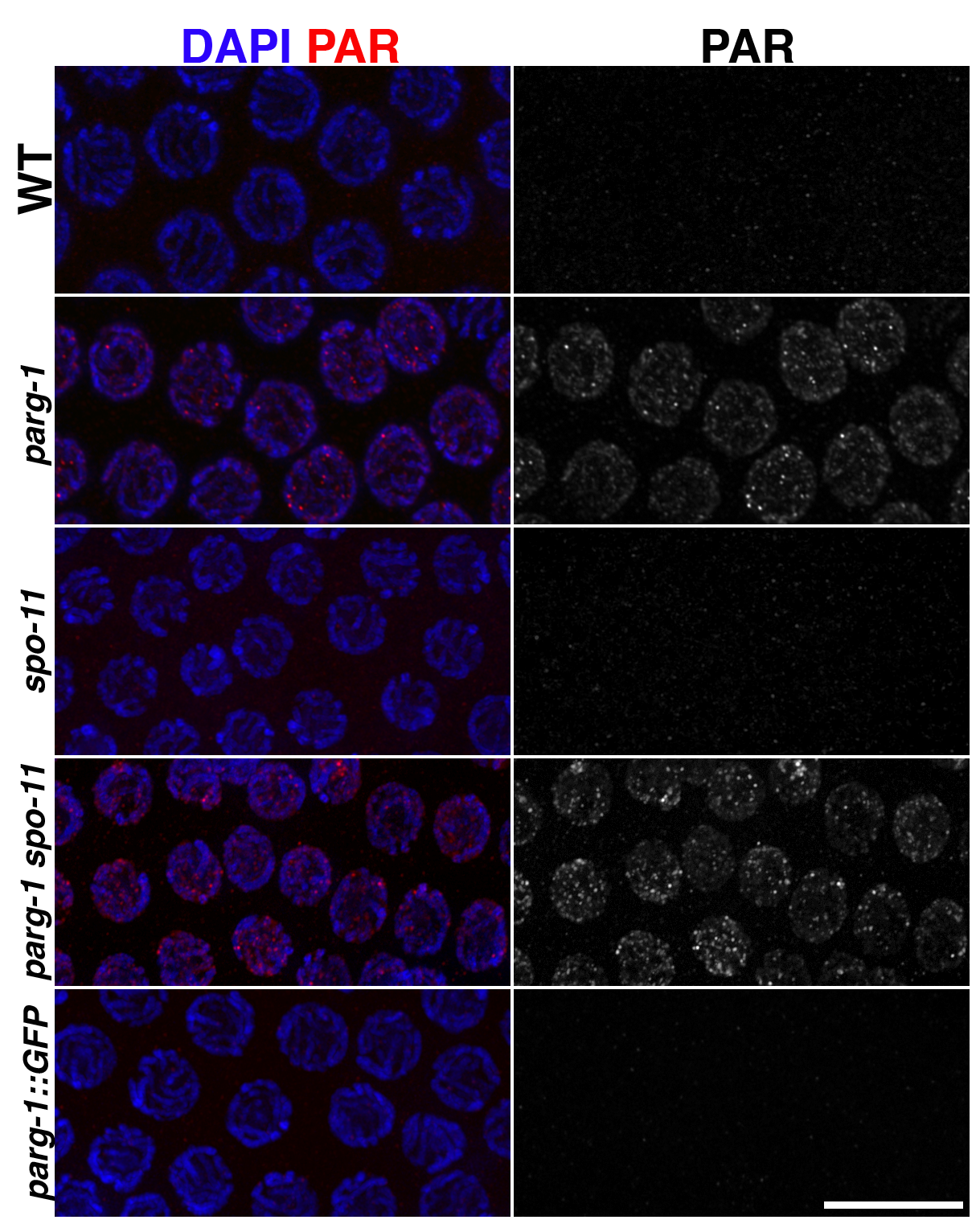

### Supplemental Fig 2

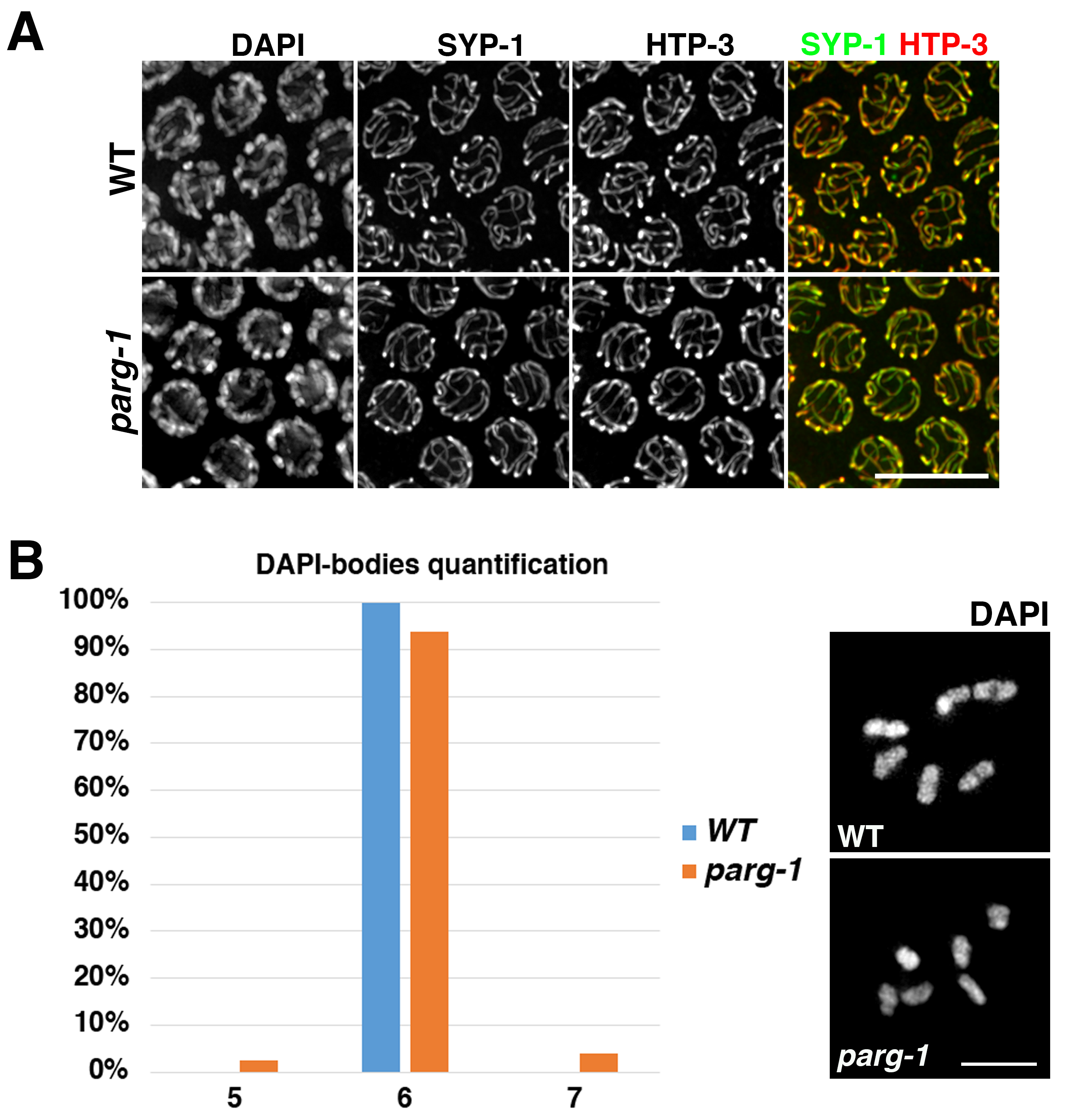

### Supplemental Fig 3

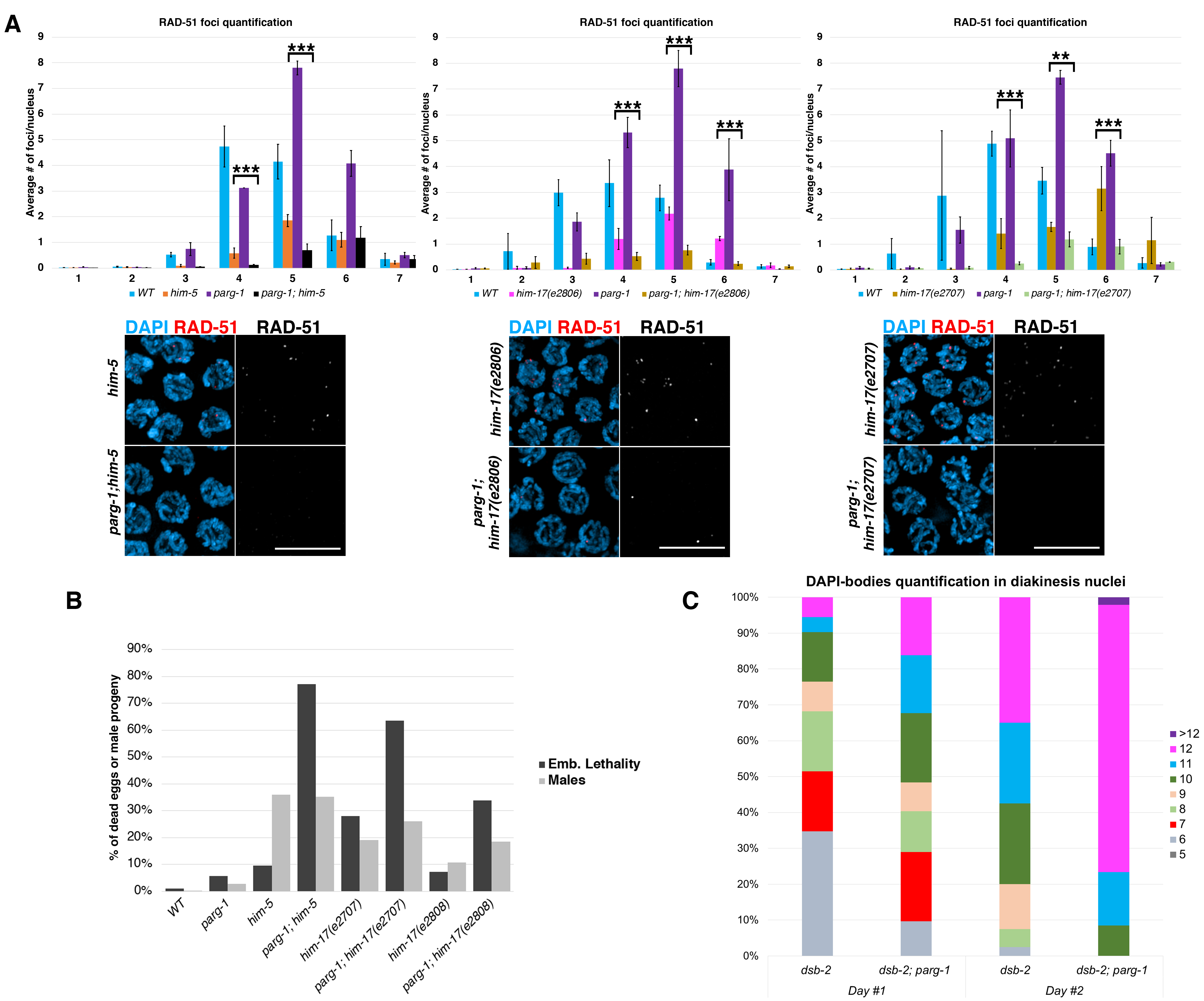

### Supplemental Fig 4

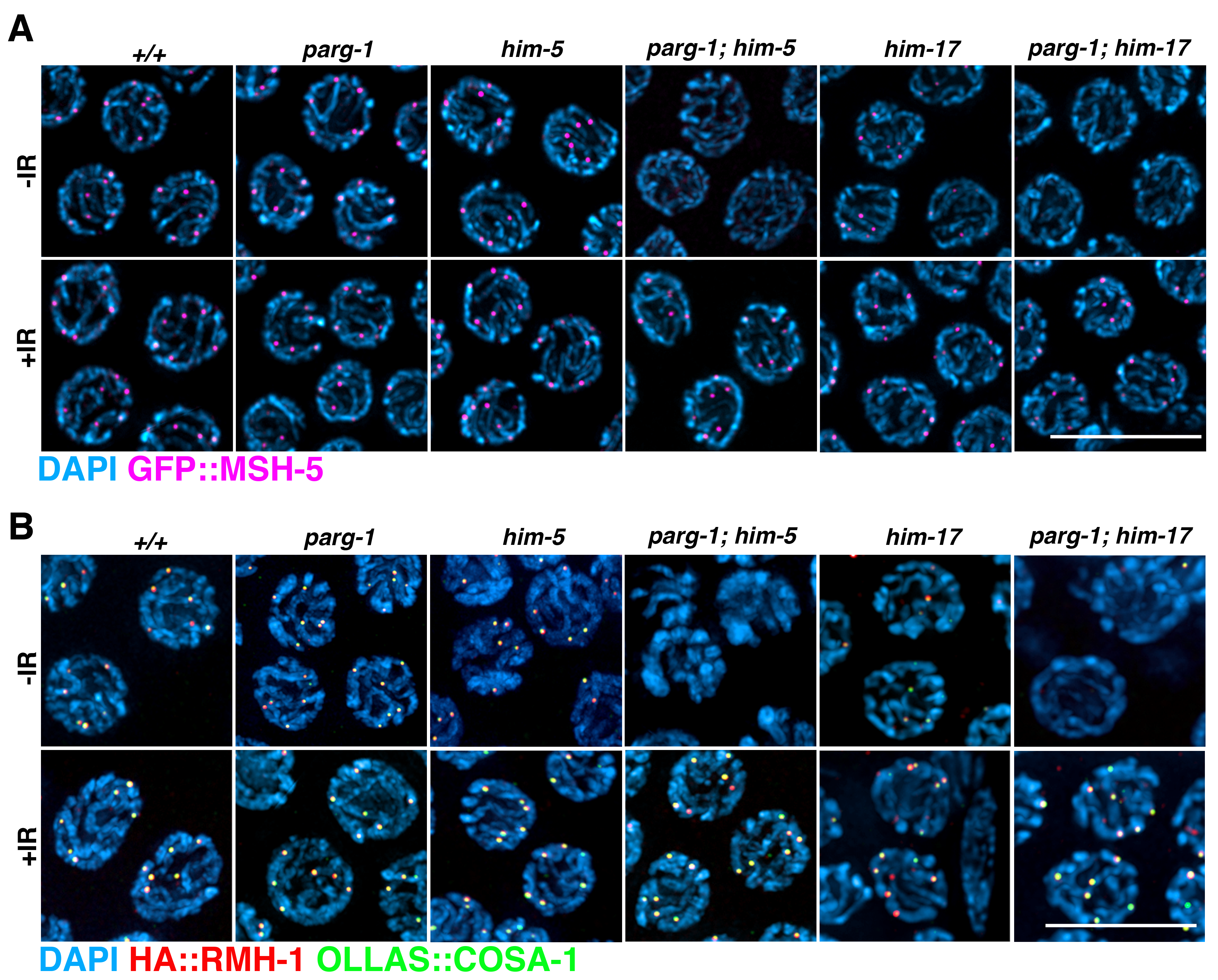

### Supplemental Fig 5

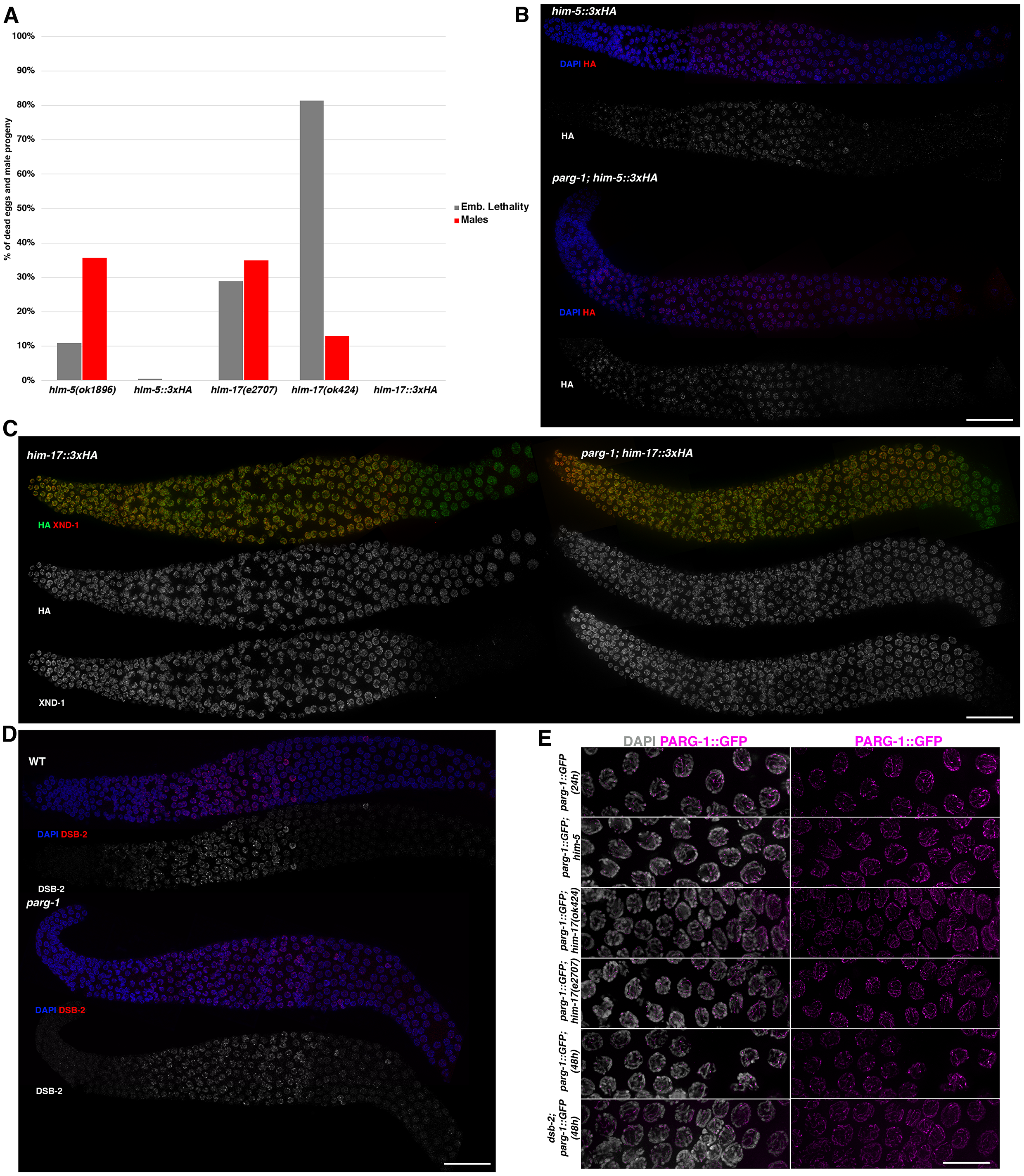

### Supplemental Fig 6

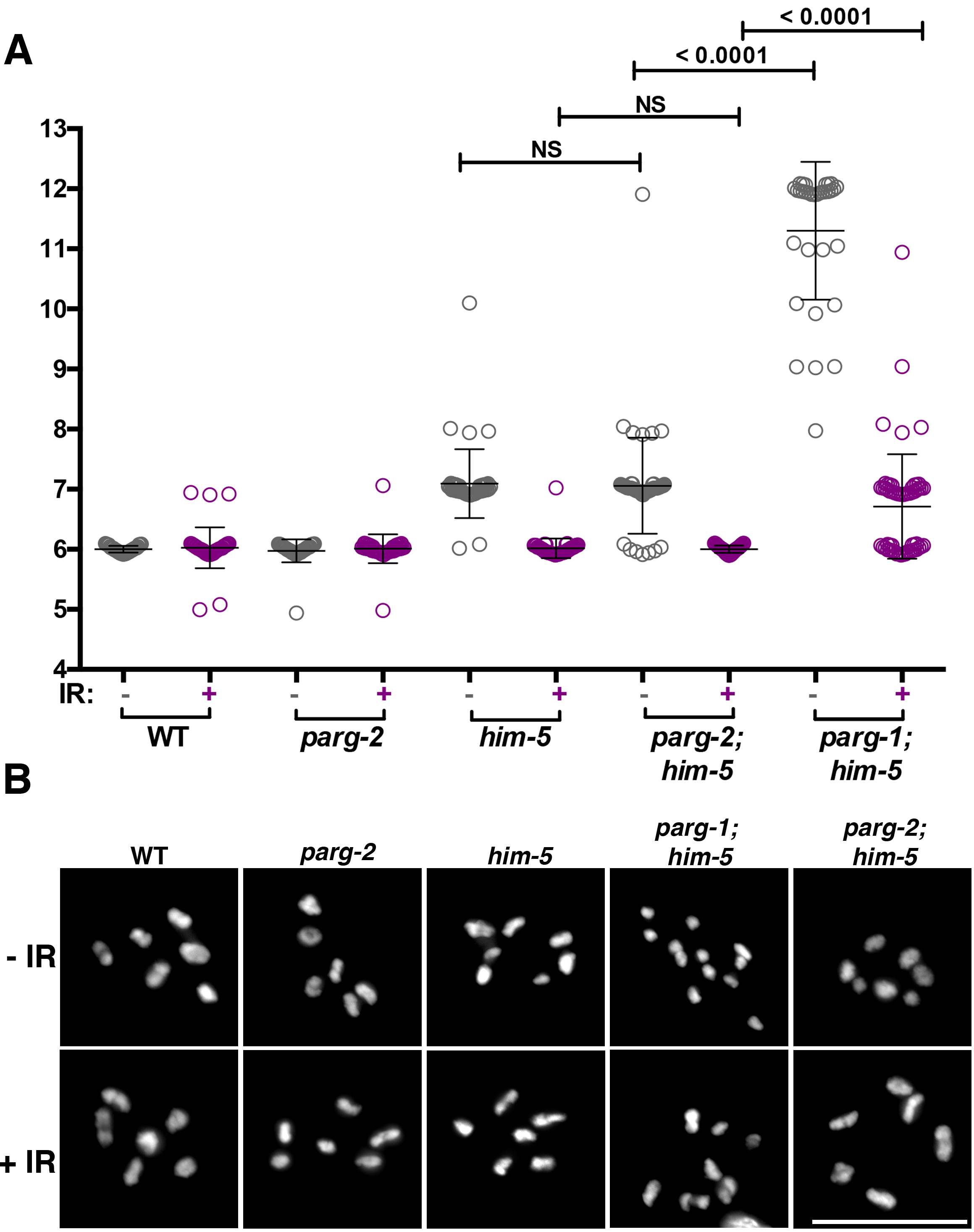

### Supplemental Fig 7

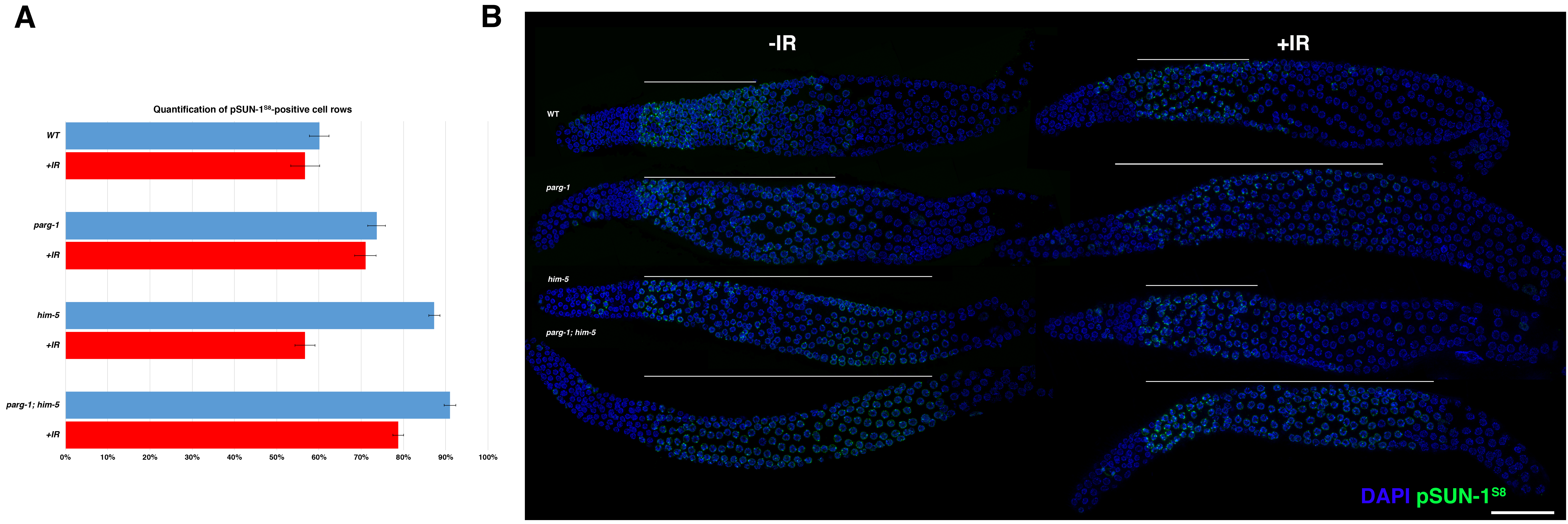

### Supplemental Fig 8

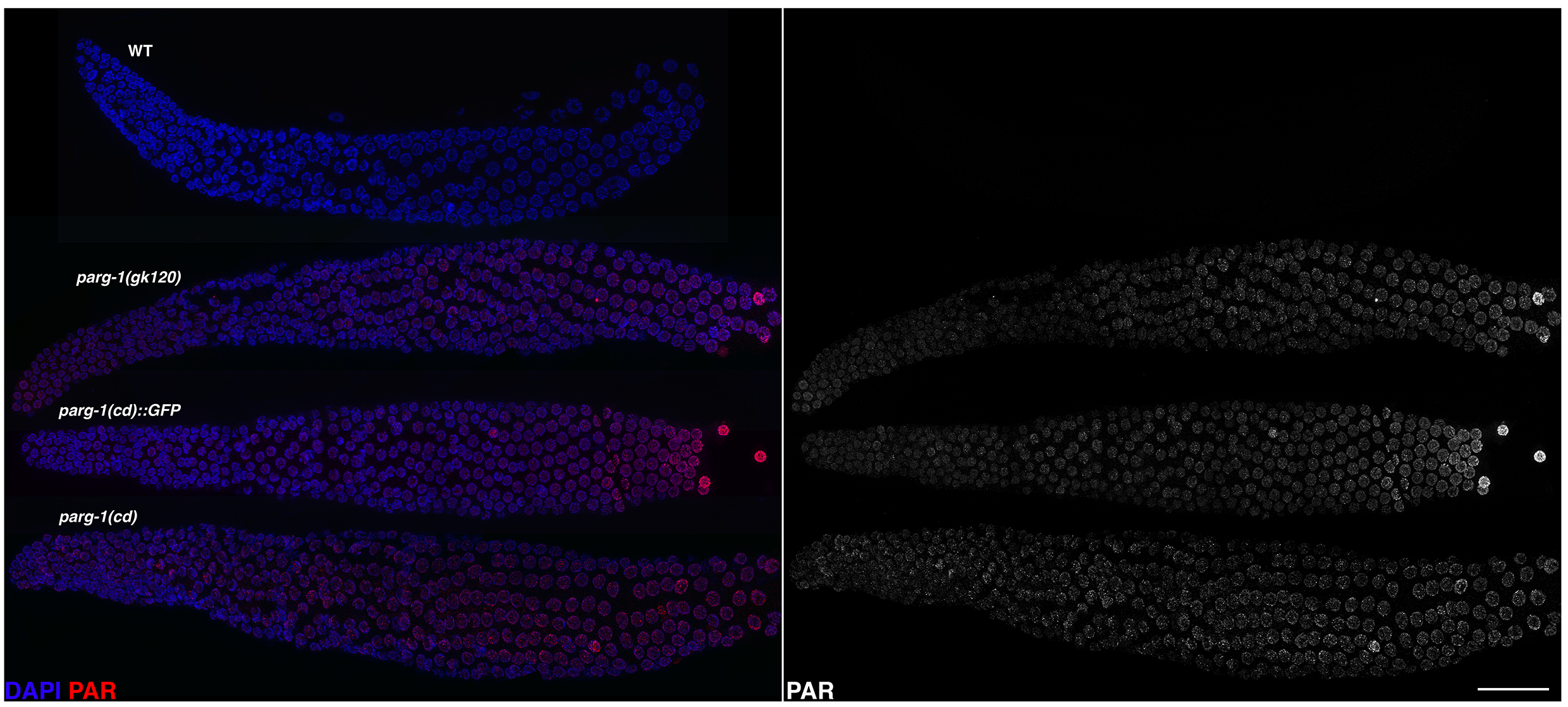
