## Supplementa Table S1 for "Poly(ADP-ribose) glycohydrolase promotes formation and homology-directed repair of meiotic DNA double-strand breaks independent of its catalytic activity"

| Genotype | Zone 1 | Zone 2 | Zone 3 | Zone 4 | Zone 5 |
|---|---|---|---|---|---|
| *GFP::msh-5* | 179 | 142 | 159 | 117 | 72 |
| *GFP::msh-5 +IR* | 130 | 170 | 212 | 165 | 105 |
| *GFP::msh-5 parg-1* | 207 | 178 | 176 | 143 | 58 |
| *GFP::msh-5 parg-1 +IR* | 234 | 194 | 211 | 129 | 74 |
| *GFP::msh-5; him-5* | 276 | 239 | 205 | 170 | 107 |
| *GFP::msh-5; him-5 +IR* | 235 | 246 | 219 | 149 | 82 |
| *GFP::msh-5 parg-1; him-5* | 195 | 166 | 141 | 114 | 79 |
| *GFP::msh-5 parg-1; him-5 +IR* | 225 | 202 | 190 | 126 | 63 |
| *GFP::msh-5; him-17* | 379 | 359 | 268 | 215 | 131 |
| *GFP::msh-5; him-17 +IR* | 181 | 178 | 162 | 100 | 76 |
| *GFP::msh-5 parg-1; him-17* | 454 | 417 | 329 | 251 | 230 |
| *GFP::msh-5 parg-1; him-17 +IR* | 182 | 147 | 134 | 84 | 54 |

Number of nuclei scored for quantification of GFP::MSH-5 foci.

| Genotype | Zone 1 | Zone 2 | Zone 3 | Zone 4 | Zone 5 |
|---|---|---|---|---|---|
| *HA::rmh-1* | 196 | 224 | 181 | 153 | 101 |
| *HA::rmh-1 +IR* | 204 | 195 | 171 | 134 | 79 |
| *HA::rmh-1; parg-1* | 209 | 198 | 148 | 130 | 106 |
| *HA::rmh-1; parg-1 +IR* | 174 | 141 | 131 | 87 | 60 |
| *HA::rmh-1; him-5* | 201 | 218 | 172 | 135 | 80 |
| *HA::rmh-1; him-5 +IR* | 201 | 123 | 118 | 78 | 50 |
| *HA::rmh-1; parg-1; him-5* | 209 | 168 | 136 | 89 | 78 |
| *HA::rmh-1; parg-1; him-5 +IR* | 226 | 175 | 141 | 102 | 59 |
| *HA::rmh-1; him-17* | 226 | 207 | 158 | 101 | 85 |
| *HA::rmh-1; him-17 +IR* | 162 | 144 | 96 | 86 | 68 |
| *HA::rmh-1; parg-1; him-17* | 189 | 155 | 138 | 99 | 92 |
| *HA::rmh-1; parg-1; him-17 +IR* | 234 | 215 | 155 | 97 | 75 |

Number of nuclei scored for quantification of HA::RMH-1 foci.

| Genotype | 0 Gy | 10 Gy +8h | 10 Gy +17h |
|---|---|---|---|
| *OLLAS::cosa-1* | 130 | 170 | 118 |
| *OLLAS::cosa-1; parg-1* | 119 | 101 | 102 |
| *OLLAS::cosa-1; him-5* | 109 | 128 | 156 |
| *OLLAS::cosa-1; parg-1; him-5* | 206 | 113 | 132 |

Number of nuclei scored for quantification of OLLAS::COSA-1 foci.

| Genotype | 0 Gy | 10 Gy +8h | 10 Gy +17h |
|---|---|---|---|
| *OLLAS::cosa-1* | 108 | 177 | 149 |
| *OLLAS::cosa-1; parg-1* | 116 | 112 | 125 |
| *OLLAS::cosa-1; him-5* | 116 | 140 | 184 |
| *OLLAS::cosa-1; parg-1; him-5* | 234 | 131 | 160 |

Number of nuclei scored for quantification of synapsis (from Fig. 5).

| **Genotype** | **0 Gy** | **10 Gy** |
|---|---|---|
| WT | 215 | 205 |
| *parg-1(cd)* | 137 | 146 |
| *him-5* | 169 | 190 |
| *parg-1(cd); him-5* | 165 | 212 |

Number of nuclei scored for quantification of synapsis (from Fig. 6).
