## Supplementa Table S2 for "Poly(ADP-ribose) glycohydrolase promotes formation and homology-directed repair of meiotic DNA double-strand breaks independent of its catalytic activity"

| Genotype | Zone 1 | Zone 2 | Zone 3 | Zone 4 | Zone 5 | Zone 6 | Zone 7 |
|---|---|---|---|---|---|---|---|
| WT | 222 | 196 | 158 | 137 | 146 | 121 | 117 |
| *parg-1* | 130 | 170 | 122 | 114 | 105 | 79 | 93 |
| *spo-11* | 239 | 233 | 218 | 199 | 156 | 155 | 93 |
| *spo-11 parg-1* | 197 | 211 | 200 | 185 | 132 | 102 | 78 |
| *him-5* | 257 | 250 | 182 | 191 | 181 | 145 | 79 |
| *parg-1; him-5* | 180 | 220 | 177 | 172 | 103 | 153 | 97 |
| *him-17(e2806)* | 273 | 233 | 238 | 255 | 254 | 201 | 124 |
| *parg-1; him-17(e2806)* | 199 | 198 | 150 | 126 | 166 | 143 | 88 |
| *him-17(e2707)* | 204 | 197 | 171 | 174 | 128 | 78 | 55 |
| *parg-1; him-17(e2707)* | 172 | 189 | 154 | 135 | 107 | 80 | 78 |

Number of nuclei scored for quantification of RAD-51 foci

| Genotype | 0 Gy | 10 Gy |
|---|---|---|
| WT | 34 | 50 |
| *parg-1* | 32 | 41 |
| *com-1* | 36 | 54 |
| *com-1; parg-1* | 41 | 47 |
| *mre-11(iow1)* | 124 | 51 |
| *parg-1; mre-11(iow1)* | 109 | 41 |

Quantification of DAPI-bodies in diakinesis nuclei (from Fig. 3)

| Genotype | n |
|---|---|
| WT | 40 |
| *parg-1(cd)* | 56 |
| *him-5* | 36 |
| *parg-1(cd); him-5* | 42 |
| *parg-1; him-5* | 44 |
| *parp-1; parg-1; him-5* | 35 |
| *parp-2; parg-1; him-5* | 35 |
| *parp-1; parp-2; parg-1; him-5* | 51 |

Quantification of DAPI-bodies in diakinesis nuclei (from Fig. 6)
