## Supplementa Table S3 for "Poly(ADP-ribose) glycohydrolase promotes formation and homology-directed repair of meiotic DNA double-strand breaks independent of its catalytic activity"

| Genotype | Source | Id. number |
|---|---|---|
| *C. elegans/ parp-1(ddr31) I.* | This study | NSV167 |
| *C. elegans/ rmh-1(jf172[HA::rmh-1]) I.* | This study | NSV240 |
| *C. elegans/ htp-3::TEV::eGFP::myc::3xFLAG I.* | CGC | JH4008 |
| *C. elegans/ parp-2(ok344) II.* | CGC | VC1171 |
| *C. elegans/ dsb-2(me96) II.* | Villeneuve lab | AV477 |
| *C. elegans/ GFP::cosa-1 II.* | Villeneuve lab | AV630 |
| *C. elegans/ cosa-1(tm3298)/qC1[dpy-19(e1259) glp-1(q339) qIs26] III.* | Villeneuve lab | AV590 |
| *C. elegans/ oxTi574 II; unc-119(ed3) III.* | CGC | EG7875 |
| *C. elegans/ cosa-1(ddr12[OLLAS::cosa-1]) III.* | Janisiw et al.; 2018 | NSV97 |
| *C. elegans/ parg-1(gk120) IV.* | CGC | VC130 |
| *C. elegans/ parg-2(ddr20) IV.* | This study | NSV140 |
| *C. elegans/ parg-1(ddr3[parg-1::GFP]) IV.* | This study | NSV25 |
| *C. elegans/ parg-1(gk120) parg-2(ddr20) IV.* | This study | NSV126 |
| *C. elegans/ msh-5(ddr22[GFP::msh-5]) IV* | Janisiw et al.; 2018 | NSV129 |
| *C. elegans/ parg-1(ddr29[parg-1*$^{E554,555A}$*::GFP]) IV.* | This study | NSV162 |
| *C. elegans/ parg-1(ddr34[parg-1*$^{E554,555A}$*]) IV.* | This study | NSV177 |
| *C. elegans/ parg-1*$^{gk120}$*(ddr51) IV in CB4856.* | This study | NSV278 |
| *C. elegans/ him-5(ok1896) V.* | CGC | RB1562 |
| *C. elegans/ him-17(e2806) V.* | This study | NSV169 |
| *C. elegans/ him-17(e2707) V.* | This study | NSV170 |
| *C. elegans/ him-17(ddr37[him-17::3xHA]) V.* | This study | NSV205 |
| *C. elegans/ syp-2(ok307)7nT1 [unc-?(n754) let-?(m435)] (IV;V).* | CGC | AV276 |
| *C. elegans/ parp-2 (ok344) II; parg-1(gk120) IV.* | This study | NSV09 |
| *C. elegans/ spo-11(ok79)/nT1 [unc-?(n754) let-?] (IV;V).* | CGC | AV106 |
| *C. elegans/ syp-3(ok758) I; ieSi11 II; unc-119(ed3) III.* | CGC | CA1218 |
| *C. elegans/ parg-1(gk120) IV; him-5(ok1896) V.* | This study | NSV56 |
| *C. elegans/ parg-1(gk120) spo-11(ok79)/nT1 [unc-?(n754) l et-?] (IV;V).* | This study | NSV74 |
| *C. elegans/ parg-1(gk120) with Hawaiian chromosome V* | This study | NSV121 |
| *C. elegans/ parg-1(ddr3[parg-1::GFP]) IV; him-5(ok1896) V.* | This study | NSV124 |
| *C. elegans/ cosa-1(ddr12[OLLAS::cosa-1]) III; parg- 1(gk120) IV.* | This study | NSV137 |
| *C. elegans/ cosa-1(ddr12[OLLAS::cosa-1]) III; him- 5(ok1896) V.* | This study | NSV138 |
| *C. elegans/ cosa-1(ddr12[OLLAS::cosa-1]) III; parg- 1(gk120) IV; him-5(ok1896) V.* | This study | NSV139 |
| *C. elegans/ cosa-1(ddr12[OLLAS::cosa-1]) III; parg- 1(gk120); him-5(ok1896)/nT1[unc-?(n754) let-?] (IV* | This study | NSV155 |
| *C. elegans/ parg-1(gk120) msh-5(ddr22[GFP::msh-5]) IV* | This study | NSV157 |
| *C. elegans/ parg-1(ddr3[parg-1::GFP]) IV; syp- 2(ok307)7nT1 [unc-?(n754) let-?(m435)] (IV;V).* | This study | NSV160 |
| *C. elegans/ dsb-2(me96) II; parg-1(gk120) IV.* | Yanowitz lab | QP1367 |
| *C. elegans/ mre-11(iow1)/nT1 [qIs51] (IV;V).* | CGC | SSM2 |
| *C. elegans/ parg-1(gk120) IV; him-17(e2806) V.* | Yanowitz lab | QP1374 |
| *C. elegans/ parg-1(gk120) IV; mre-11(iow1)/nT1 [qIs51] (IV;V).* | Yanowitz lab | QP1377 |
| *C. elegans/ parp-1(ddr31) I; parp-2(ok344) II.* | This study | NSV175 |

| | | |
|---|---|---|
| C. elegans/ parg-1(ddr34[parg-1$^{E554,555A}$]) IV in CB4856. | This study | NSV178 |
| C. elegans/ rmh-1(jf54)/ hT2 [bli-4(e937) let-?(q782) qIs48] (I;III). | This study | NSV180 |
| C. elegans/ parg-1(ddr34[parg-1$^{E554,555A}$::GFP]) IV; him- 5(ok1896) V. | This study | NSV181 |
| C. elegans/ com-1(t1626) unc-32(e189)/hT2 [bli-4(e937) let- ?(q782) qIs48] (I;III). | This study | NSV182 |
| C. elegans/ parp-1(ddr31) I; parg-1(gk120) IV. | This study | NSV184 |
| C. elegans/ cosa-1(tm3298)/qC1[dpy-19(e1259) glp- 1(q339) qIs26] III; parg-1(ddr3[parg-1::GFP]) IV. | This study | NSV186 |
| C. elegans/ parg-1(ddr34[parg-1$^{E554,555A}$]) IV; him-5(ok1896) V. | This study | NSV190 |
| C. elegans/ com-1(t1626) unc-32(e189)/hT2 [bli-4(e937) let- ?(q782) qIs48] (I;III); parg-1(gk120) IV. | This study | NSV194 |
| C. elegans/ him-17(ok424)/nT1 [unc-?(n754) let-?] (IV;V). | This study | NSV196 |
| C. elegans/ parp-1(ddr31) I; parp-2(ok344) II; parg-1(gk120) IV. | This study | NSV199 |
| C. elegans/ parg-1(gk120) IV; him-17(ddr37[him-17::3xHA] V. | This study | NSV206 |
| C. elegans/ parg-1(gk120)/nT1 [unc-?(n754) let-?] (IV;V).; him-17(ok424)/nT1 [unc-?(n754) let-?] (IV;V). | This study | NSV209 |
| C. elegans/ parg-1(gk120)/nT1 [unc-?(n754) let-?] (IV;V).; him-17(e2707)/nT1 [unc-?(n754) let-?] (IV;V). | This study | NSV236 |
| C. elegans/ msh-5(ddr22[GFP::msh-5]) IV; him-5(ok1896) V. | This study | NSV241 |
| C. elegans/ msh-5(ddr22[GFP::msh-5]) IV; him-17(e2707) V. | This study | NSV242 |
| C. elegans/ parp-2(ok344) II; parg-1(gk120) IV; him- 5(ok1896) V. | This study | NSV244 |
| C. elegans/ him-17(e2707)/nT1 [unc-?(n754) let-?] (IV;V). | This study | NSV246 |
| C. elegans/ msh-5(ddr22[GFP::msh-5])/nT1 [unc-?(n754) l et-?] (IV;V); him-17(e2707)/nT1 [unc-?(n754) let-?] (IV;V). | This study | NSV247 |
| C. elegans/ him-5(ddr43[him-5::3xHA]) V. | This study | NSV250 |
| C. elegans/ parg-1(gk120) IV; him-5(ddr43[him-5::3xHA]) V. | This study | NSV252 |
| C. elegans/ rmh-1(jf172[HA::rmh-1]); cosa- 1(ddr12[OLLAS::cosa-1]) III; parg-1(gk120) IV. | This study | NSV253 |
| C. elegans/ parg-1(gk120) msh-5(ddr22[GFP::msh-5])/nT1 [unc-?(n754) let-?] (IV;V); him-17(e2707)/nT1 [unc-?(n754) let-?] (IV;V). | This study | NSV254 |
| C. elegans/ htp-3(tm3655)/hT2 [bli-4(e937) let-?(q782) ql (I;III). | This study | NSV255 |
| C. elegans/ rmh-1(jf172[HA::rmh-1]); cosa- 1(ddr12[OLLAS::cosa-1]) III; him-5(ok1896) V. | This study | NSV256 |
| C. elegans/ rmh-1(jf172[HA::rmh-1]); cosa-1(ddr12[OLLAS::cosa-1]) III; parg-1(gk120) IV; him- 5(ok1896) V. | This study | NSV257 |
| C. elegans/ rmh-1(jf172[HA::rmh-1]) I; cosa- 1(ddr12[OLLAS::cosa-1]) III. | This study | NSV258 |
| C. elegans/ parg-1(ddr3[parg-1::GFP]) IV; him-5(ddr43[him- 5::3xHA]) V. | This study | NSV262 |

| | | |
|---|---|---|
| *C. elegans/ htp-3(tm3655)/hT2 [bli-4(e937) let-?(q782) qI (I;III); parg-1(ddr3[parg-1::GFP]) IV.* | This study | NSV265 |
| *C. elegans/ parg-1(ddr3[parg-1::GFP]) IV; him- 17(ddr37[him-17::3xHA]) V.* | This study | NSV267 |
| *C. elegans/ parg-1(gk120) msh-5(ddr22[GFP::msh-5]) IV; him-5(ok1896) V.* | This study | NSV273 |
| *C. elegans/ rmh-1(jf172[HA::rmh-1])I; cosa- 1(ddr12[OLLAS::cosa-1]) III; him-17(e2707) V.* | This study | NSV276 |
| *C. elegans/ dsb-2(me96) II; parg-1(ddr3[parg-1::GFP]) IV.* | This study | NSV280 |
| *C. elegans/ parg-1(ddr3[parg-1::GFP]) IV; him-17(e2707) V.* | This study | NSV281 |
| *C. elegans/ rmh-1(jf172[HA::rmh-1]) I; cosa- 1(ddr12[OLLAS::cosa-1]) III; parg-1(gk120)/nT1 [unc- ?(n754) let-?] (IV;V); him-17(e2707)/nT1 [unc-?(n754) let-?] (IV;V).* | This study | NSV286 |
| *C. elegans/ cosa-1(ddr12[OLLAS::cosa-1]) III; parg- 1(ddr3[parg-1::GFP]) IV.* | This study | NSV292 |
| *C. elegans/ parg-1(ddr3[parg-1::GFP])/nT1 [unc-?(n754) l et-?] (IV;V); him-17(ok424)/nT1 [unc-?(n754) let-?] (IV;V).* | This study | NSV294 |
